# A Mimicry-Based Strategy Between Human and Commensal Antigens for the Development of a New Family of Immune Therapies for Cancer

**DOI:** 10.1101/2024.05.27.596102

**Authors:** Alice Talpin, Ana Maia, Jean-Marie Carpier, Guillaume Kulakowski, Camille Gaal, Francesco Strozzi, Coline Billerey, Lucie Aubergeon, Ludivine Amable, Jérôme Kervevan, Tifanny Mersceman, Alexandrine Garnier, Catia Pereira Oliveira, Carolina Calderon, Diana Bachrouche, Chloé Ventujol, Jennifer Martinez, Michaël Bonnet, Julie Noguerol, Karl Laviolette, Laura Boullerot, Marine Malfroy, Grégoire Chevalier, Olivier Adotevi, Olivier Joffre, Ahmed Idbaih, Maria Vieito, François Ghiringhelli, Agostina Stradella, Ghazaleh Tabatabai, Michael C. Burger, Iris Mildenberger, Ulrich Herrlinger, David A. Reardon, Wolfgang Wick, Cécile Gouttefangeas, Christophe Bonny, Laurent Chêne, Joao Gamelas Magalhaes

## Abstract

Peptide vaccines have emerged as a promising strategy for cancer immunotherapy, yet often lack of strong, specific and sustained immune responses against tumor antigens. To achieve a robust immune response, the effective selection of tumour antigens is crucial. While neoantigens trigger potent immune responses, their use suffers from patient specificity and their rarity in low-mutational tumors. Alternatively, the immunogenic potential of tumor-associated antigens (TAAs) is limited by central immune tolerance. Molecular mimicry and T cell cross-reactivity is a proposed mechanism to trigger a robust T cell-mediated antitumor response. Although molecular mimicry between pathogens and tumor antigens has been described, the potential benefits of exploiting this molecular mimicry with commensal bacterial antigens in antitumor immunity have not been thoroughly investigated despite strong evidence that the composition of the human microbiota significantly influences immune competency. Our new approach called OncoMimics™, which uses molecular mimicry between commensal bacterial and tumoral antigens to induce cross-reactive cytotoxic T cells against tumor cells. In preclinical studies, vaccination with OncoMimic™ peptides (OMPs) led to the expansion of CD8^+^ T cells reacting against homologous tumor-associated antigen peptides and elicits cytotoxic activity against tumor cells. OMPs are efficiently recognized by a prevalent T cell population within the peripheral blood mononuclear cells of healthy individuals. An ongoing clinical trial (NCT04116658) using OncoMimics™ in patients with glioblastoma demonstrates early, durable, and cross-reactive tumor antigen CD8^+^ T cell responses with pronounced memory persistence. By overcoming the current vaccine limitations, OncoMimics™ constitutes a promising strategy for enhancing cancer immunity and improving patient outcomes.

**Statement of Significance:** This study introduces OncoMimics™, a peptide-based immunotherapy leveraging molecular mimicry to induce robust, cross-reactive T cell responses against tumor antigens, showing promising early results in an ongoing glioblastoma clinical trial (NCT04116658)

## Introduction

Immune checkpoint inhibitors (ICIs) offer a potential curative cancer treatment by boosting anti-cancer T cell immune responses in treated patients. However, their inability to stimulate specific T cells in ‘cold’ tumors significantly limits their efficacy. Therapeutic vaccines could ideally complement ICI treatments in cancers with low mutational burdens and limited spontaneous T cell responses (1).

Despite the potential of therapeutic vaccines, most vaccine trials have failed, largely due to their inability to sustainably stimulate tumor-reacting T cells (2). Effective induction of anti-tumor cytotoxic T lymphocyte (CTL) responses critically depends on selecting appropriate tumor antigens. Typically, tumors present tumor-associated antigens (TAAs) shared among patients, offering broad applicability yet limited immunogenicity. This limitation stems from thymic negative selection, which eliminates highly self-reactive T cells to prevent autoimmunity, thereby reducing the affinities of T-cell receptor (TCR) repertoire for these antigens. Neoantigens, not subject to thymic deletion, elicit stronger immunogenicity; however, their clinical applicability is limited due to patient specificity and rarity in low mutational burden tumors (3,4). We have developed a novel approach to enhance the immunogenicity of natural TAAs, enabling the human immune system to activate endogenous T cells that recognize key tumor drivers and achieve tumor shrinkage, even in cold tumors such as glioblastoma. We leveraged two well-described mechanisms to enhance CD8^+^ T cell recognition of tumors: peptide molecular mimicry, which exploits structural similarities between tumor and microbial peptides to engage pre-existing T cell responses, and TCR cross-reactivity, enabling a single TCR to recognize diverse antigens through flexible binding capabilities (5,6).

We have developed a systematic bioinformatic approach to identify similarities between bacterial antigens and human TAAs. The microbiome encodes billions of “foreign” antigens that can potentially trigger high-affinity TCRs preserved from central immune tolerance. Furthermore, the composition of the gut microbiome impacts ICI therapy outcome (7–9). Thus, exposure to commensal epitopes might generate cross-reactive T cells that could be leveraged to maximize the efficacy of peptide-based immunotherapies (10). Taking advantage of the widespread presence of these bacteria in the human population, we expected to stimulate a diverse range of memory T cells capable of cross-reacting with TAAs. Furthermore, by employing commensal-derived-peptides (CDPs) with high affinity for HLA molecules, we effectively address the limited immunogenicity of tumor-associated antigen derived peptides (TAAps) and aim to reactivate a robust memory T cell repertoire that is primarily maintained to protect us against commensal bacteria.

We describe here this approach, termed ‘oncomimicry,’ which enables the discovery of commensal-derived short peptides mimicking TAAps, eliciting potent CTL responses. The selection process for these OncoMimic™ peptides (OMPs) relies on various criteria, including sequence homologies between OMPs and their TAA counterparts, binding affinities to HLA class I allelic products, predicted cleavage scores, and frequencies of commensal bacteria sources expressing the selected OMPs in the human population. OMP candidates meeting these criteria were tested and validated for their immunogenicity and ability to elicit cross-reacting TAA-specific CTL responses in HLA-A2-humanized mice, resulting in tumor regression. Ex vivo experiments showed that identified OMPs stimulated human T cell proliferation and triggered cytolytic activity against target cells presenting homologous TAAs. Finally, initial data from an ongoing clinical trial (NCT04116658) demonstrated that OMPs generate fast, potent, and long-lasting immune responses in patients, providing a strong rationale for using commensal-derived peptides to enhance peptide-based immunotherapies.

## Material and Methods

### Compounds and antibodies

All synthetic peptides (>80% purity) were synthesized using sb-PEPTIDE (**Table 1**). For flow cytometry mouse phenotyping, the following monoclonal antibodies (mAbs) and compounds were used: DAPI (SIGMA), PerCP-Vio700 CD45 (clone REA737, Miltenyi Biotec), APC-Cy7 TCR-β (clone H57-597, BioLegend), FITC CD11c (clone REA754, Miltenyi Biotec), VioGreen CD11b (clone REA592, Miltenyi Biotec), and PE HLA-A2 (clone REA517, Miltenyi Biotec). The healthy volunteer peripheral blood mononuclear cells (PBMCs) fraction was phenotyped using an 8-Color Immunophenotyping Kit, anti-human, REAfinity™ (Miltenyi Biotec). For CD137 and CD8 enriched fractions the following mAbs from Miltenyi Biotec were used: VioBright-FITC CD137 (REA765), VioGreen CD45 (REA747), PE or PerCPVio700 CD4 (REA623), PerCPVio700 or VioGreen CD8 (REA734), APCVio770 CD3 (REA613), and APC CD19 (REA675). To exclude dead cells, LIVE/DEAD fixable violet or near-IR dead cell stain (Thermo Fisher) was added. For tetramer staining, PerCPVio700 or VioGreen CD8 (Miltenyi Biotec, REA734) was used. For HLA-A*02:01 tetramer staining, monomers folded with the OMP- and TAA-derived peptides were produced at Tübingen University using the classical refolding method and quality control as previously shown (11), as detailed in the tetramer production section below. Peptide-MHC monomers were tetramerized by incubation with streptavidin-PE (BioLegend and Invitrogen), -BV421, -BV785 (BD Biosciences), -APC, and -AlexaFluor488 (Life Technologies) and were subsequently stored as aliquots at −80°C according to Hadrup *et al.* (12). PE and APC-conjugated HLA-A*02:01 MHC dextramers (Immudex) were also used.

**Table 1.**
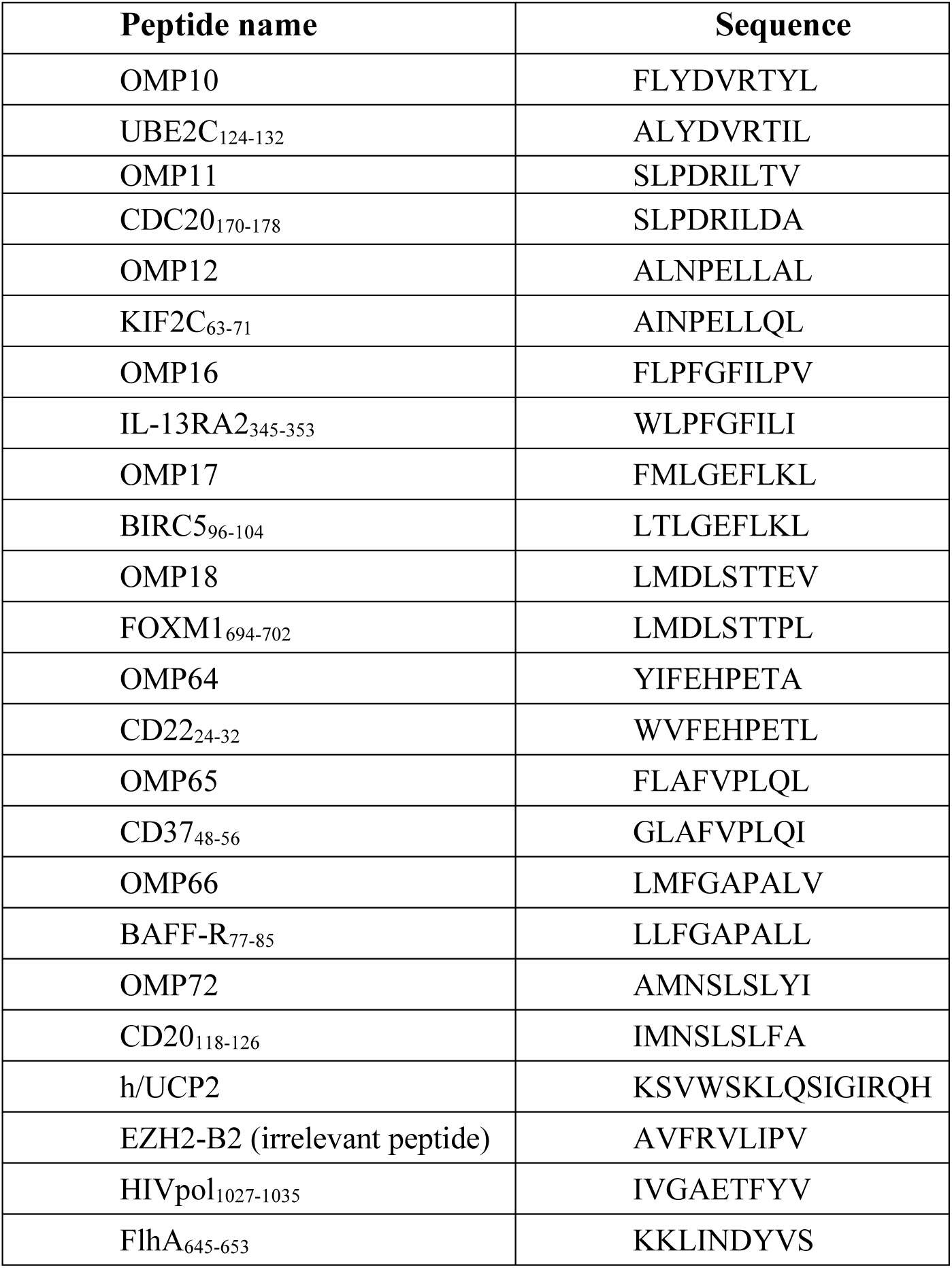
Peptide list.

### Flow cytometry

Mouse single-cell suspensions were resuspended in FACS buffer (1× PBS containing 0.5% BSA, 2mM EDTA) and incubated with Mouse BD Fc Block purified anti-mouse CD16/CD32 mAb 2.4G2 (BD Biosciences) to block non-specific staining. The cells were subsequently incubated with antibodies prepared in the FACS buffer for 30 min at 4°C. Dead cells were labelled with DAPI (Sigma) following the manufacturer’s instructions. Healthy donor (HD) PBMCs and CD137^+^ or CD8^+^-enriched cells were resuspended in FACS buffer and incubated with fluorescent-conjugated antibodies for 20 min at 4°C. Dead cells were excluded using DAPI (Sigma) or the LIVE/DEAD^TM^ Fixable Violet Dead Cell Stain Kit (Thermo Fisher Scientific) according to the manufacturer’s recommendations. After staining, the cells were washed twice with FACS buffer and analyzed using flow cytometry (MacsQuant Analyzer 10 flow cytometer, Miltenyi Biotec). The FlowJo™ software (BD Biosciences) was used to analyze the results. Data were collected from approximately 10^5^ cells. All the mAbs used for flow cytometric analysis are listed above. HLA-A*02:01-tetramer staining was performed using fluorescent-labeled tetramers (10ml/1-3×10^6^ cells for dextramers (Immudex) or 2.5ug/mL for tetramers produced at Tübingen University; the cells were incubated at room temperature (RT) for 10 min or 30 min, respectively, followed by 20 min incubation at 4°C with VioGreen CD8 mAb (REA734). Dead cells were excluded using DAPI, LIVE/DEAD^TM^ Fixable Violet, or Near-IR Dead Cell Stain Kit (Thermo Fisher Scientific).

### Cell lines and culture

The Human TAP-deficient T2 cell line (ATCC Cat# CRL-1992) was used for HLA-A*02:01 binding and stability assays of candidate peptides and for *in vitro* stimulation rounds of HD PBMCs and killing assays (13). T2 cells were cultured in Iscove’s Modified Dulbecco’s medium (IMDM) complete medium, according to the manufacturer’s protocol. The sarcoma tumor cell line (SARC-A2) was derived from a spontaneously arising sarcoma in HLA-A2/DR1 mice (14). SARC-A2 cells were transduced with a lentiviral vector encoding GFP only (SARC-A2-GFP) or carrying the CD20 full human gene upstream of an IRES element followed by a GFP cassette (SARC-A2-hCD20-GFP). A puromycin N-acetyltransferase gene was also present in the expression cassette to allow selection of transduced cells with puromycin (10μg/mL). Transduction efficiency was assessed using flow cytometry based on the expression of eGFP and human CD20 (Miltenyi Biotec).

### Measurement of peptide relative binding affinity for HLA-A*02:01

Peptide relative binding affinities for HLA-A*02:01 were measured as previously described with some minor modifications (15). Briefly, T2 cells were incubated with a titration of each tested peptide and 100 ng/mL human β2-microglobulin (Sigma) in serum-free medium (TexMacs) at 37°C for 20h. After incubation, the cells were washed and surface-stained with HLA-A2 mAb (REA517). HIVpol_1027-1035_ and FlhA_645-653_ peptides were used as positive and negative reference controls, respectively, as previously described (16). The geometric mean fluorescence intensities of HLA-A2 staining were measured by flow cytometry (MACSQuant analyzers, Miltenyi Biotec) to determine the percentage of HLA-A*02:01 molecule stabilized at the cell surface, as follows:

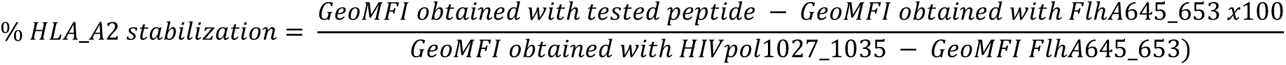

The relative affinity (RA) of each peptide for HLA-A*02:01 molecules was determined as the ratio of the peptide concentration to the reference peptide concentration (HIV) that stabilized 20% of the HLA-A2 signal at the cell surface. The lower the RA is, the stronger is the peptide binding to HLA-A*02:01.

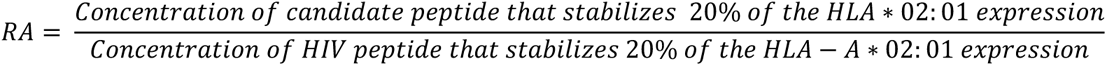

### Assessment of peptide/HLA-A*02:01 complex stability

The stability of peptide binding on HLA-A*02:01 was measured as described previously with minor modifications (15). T2 cells were pulsed overnight at 37°C with 100 μM of individual peptides in serum-free medium (TexMacs) supplemented with 100 ng/µL of human β2-microglobulin (Sigma), followed by the indicated chase kinetics performed in the presence of Brefeldin A (Sigma) to inhibit the neo-synthesis of HLA-A*02:01. At the end of the chase period, cells were surface-stained with HLA-A2 mAb (REA517) at 4°C, fixed with 0.5% paraformaldehyde, and analyzed as described for the binding assay. The dissociation complex at 50% (DC50) was defined as the half-life of the peptide-MHC complex and was calculated as the time required to observe 50% decay of the initially stabilized peptide-MHC complex.

### Animal models

All experimental protocols were approved by the local Ethics Committee on Animal Experimentation and followed the guidelines of the European Union. The previously described HLA-DRB1*01:01/HLA-A*02:01-transgenic mice (A2/DR1 mice) were obtained from Francois Lemonnier (Institut Pasteur, Paris, France) and are H-2 class I and II knockout animals (17). Their CD8^+^ and CD4^+^ T cells are restricted to HLA-A*02:01 and HLA-DR1*01:01, respectively. All the mice used in the described studies were bred and maintained under specific pathogen-free conditions at Charles River Laboratories.

### A2/DR1 prime-boost immunizations

Mice were immunized on days 0 (prime) and 14 (boost), with the indicated MHC class I peptides (5, 30, or 95 nmol per mouse) and MHC class II helper peptide (UCP2, 30-100 µg per mouse) emulsified in Montanide^TM^ ISA 51 VG (ratio 50:50, Seppic). The peptide formulation was freshly prepared, and all immunizations were performed on loose skin over the neck or subcutaneously in the flank of the animals with 100 µL of the emulsified preparation.

### Restimulation of peptide-specific CD8^+^ mouse T cells *ex vivo* post immunization

To determine the immunogenicity of the analyzed peptides and the cross-reactivity of specific T cells against human TAA homologs, mice were euthanized 21 days post-prime immunization, and the number of IFN-γ peptide-specific CD8^+^ T cells in the spleen of the animals was determined by ELISpot (Mabtech, 3321-4APT-2 kit) following the manufacturer’s recommendations. Briefly, spleens were harvested and mechanically disrupted, and red blood cells were lysed with red blood cell (RBC) lysis buffer (Miltenyi Biotec). 2×10^5^ splenocytes were restimulated per ELISpot well using the indicated peptides at a concentration of 10 µM. PMA (Phorbol-myristate acetate, 0.1µM, Sigma) and ionomycin (1µM, Sigma) were used as positive controls, and EZH2-B2, an HLA-A2:01-restricted peptide, was used as a negative control. Cells were cultured for approximately 20 h in media (RPMI-1640 (Sigma), 10% FBS (VWR), 1% GlutaMAX (Gibco), 1% non-essential amino acid (Sigma), 10mM HEPES (Sigma), 1mM sodium pyruvate (Sigma)+1% penicillin/streptomycin (Sigma), 50µM β-mercaptoethanol (Sigma), and respective peptides at 37°C and 5% CO_2_. IFN-γ spots were detected according to the manufacturer’s instructions and counted using the iSpot ELISpot Fluorospot reader system (AID). The number of spots obtained for each mouse and each condition was subtracted from the background values (cells cultured in media only) and normalized to the frequency of total T cells from each mouse, resulting in the number of specific IFN-γ-producing T cells/10^6^ T cells.

### Tumor regression model

After a prime-boost immunization, the A2/DR1 mice were subcutaneously injected 21 days post-prime immunization in the right flank with 0.5×10^6^ SARC-A2-hCD20-GFP or SARC-A2-GFP sarcoma cells resuspended in PBS. Tumor growth was evaluated twice a week using a caliper, and the mice were euthanized when their tumors reached a volume of ≥ 300 mm^2^.

### *In vivo* cytotoxicity

To test the cytotoxic function of CD8^+^ T cells that were activated following vaccination, immunized A2/DR1 mice were challenged 6 days post-boost immunization with syngeneic splenocytes pulsed with either a mix of peptides or individual peptides. To do so, unimmunized A2/DR1 donor mice were euthanized and a suspension of syngeneic splenocytes was prepared. To assess the cytotoxic response against peptide pools, splenocytes were split into 2 fractions and labeled with a cell tracking dye (CFSE or Cell Trace Violet, ThermoFisher Scientific) using a low (0.3 µM) or a high concentration (3 µM). To determine the individual contribution of each peptide from an immunization mix, the donor cell suspension was split into as many populations as needed to have control and target cells for each peptide for analysis and labeling with cell tracking dye with different fluorescence properties (Cell Trace Blue, Cell Trace Violet, CFSE, Cell Trace Yellow, Cell Trace Far Red, BioLegend, and ThermoFisher Scientific, **Figure S3A**), allowing the individual tracking of each population. The bright populations used as target cells were pulsed for 2h at 37°C with a peptide pool or a specific peptide at a final concentration of 100µM before being mixed at an equal ratio with the other populations. Cells were injected intravenously into immunized and naïve mice and the *in vivo* cytotoxic response was assessed 20h post-injection. Antigen-specific lysis was calculated as follows:

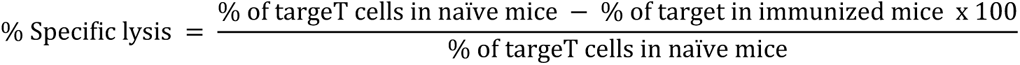

### Blood donors and isolation of PBMCs

Buffy coats from HDs were provided by the Blood Bank (Etablissement Français du Sang (EFS)) of Rungis. This study was performed in accordance with established ethical guidelines, and all blood donors provided informed consent. All blood samples were tested for HLA-A*02 (mAb REA517, Miltenyi Biotec), and HLA-A*02-positive PBMCs were isolated using Ficoll density gradient separation (GE Healthcare).

### Generation of Antigen (Ag)-specific CD8^+^ cytotoxic T cells in HLA-A*0201 HD PBMCs

PBMCs were stimulated for 24h with OMPs at 10µM in ImmunoCult^™^ medium (StemCell). For enrichment of activated Ag-specific T cells, cells were separated using CD137 magnetic bead-based positive selection as previously described (18). In brief, cells were stained with an CD137-VioBright-FITC mAb (REA765, Miltenyi Biotec) and subsequently with anti-FITC microbeads and enriched by magnetic isolation using LS MACS columns (Miltenyi Biotec). After CD137 enrichment, polyclonal T cell expansion was performed for 8 days with ImmunoCult™ Human CD3/CD28 T Cell Activator (StemCell) in ImmunoCult™ culture medium supplemented with IL-2 (50U/mL), IL-7 (12,5ng/mL), IL-15 (12,5ng/mL), and IL-21 (62,5ng/mL) (all from Miltenyi Biotec). For the next rounds of expansion (every 10days), antigen-specific CD8^+^ T cells were co-cultured with peptide-loaded T2 cells at a ratio of 10:1 in ImmunoCult ^TM^ medium supplemented with all cytokines described above. Before the co-culture, T2 cells were treated for 1h with Mitomycin C (Sigma) at 20µg/ml (19). Mitomycin C treated T2 cells were loaded with 100 ng/mL β2-microglobulin (Sigma) and 10µM OMPs overnight. These *in vitro* stimulation (IVS) rounds were repeated up to four times, as described above. The frequency of antigen-specific CD8^+^ T cells was assessed by surface staining with CD8 VioGreen mAb (REA734) and fluorescently labeled tetramers. The percentage cross-reactivity of each healthy donor (%) was calculated as follows:

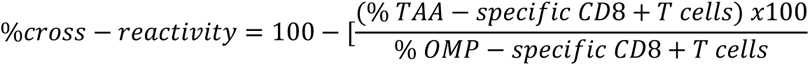

### Flow cytometry-based cytotoxicity assay

T2 target cells were labeled with CellTrace^TM^ Far Red Proliferation Kit (Thermo Fisher Scientific) and loaded overnight with 10µM individual peptide and 100ng/mL β2-microglobulin. Peptide-loaded target cells were co-cultured with peptide-specific CD8^+^ T cells at 37°C for 24h (20). Media and EZH2-B2 irrelevant peptide were used as negative controls. Antigen-specific CD8^+^ T cells were stained using VioGreen or PerCPVio700 CD8 mAbs (REA734), and dead cells were identified using the LIVE/DEAD^TM^ Fixable Violet Dead Cell Stain Kit. Conditions were performed in duplicate, and standard deviations were calculated. The percentage of specific cell lysis was calculated as follows:

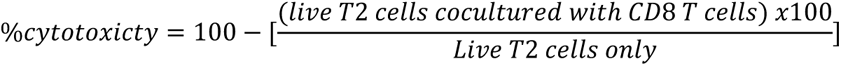

Statistical analysis was performed using GraphPad Prism (version 8).

### Trial design and treatment

An ongoing multicenter phase 1/2 trial (NCT04116658) investigated EO2401 (300 µg/peptide, every 2 weeks × 4, then every 4 weeks) and EO2401 with nivolumab (3 mg/kg every 2 weeks) in consenting patients (HLA-A2, KPS ≥70, dexamethasone ≤ 2 mg/day within 14 days before the study, normal organ function, no contraindications) with glioblastoma at first progression/recurrence after surgery and adjuvant radiotherapy/temozolomide. Treatment is administered until toxicity or tumor progression using the iRANO criteria (21). EO2401 is a therapeutic peptide vaccine composed of three HLA-A*02:01 commensal-derived peptides mimicking cytotoxic CD8^+^ T cell epitopes from the TAAs interleukin-13 receptor alpha-2 (IL-13RA2), baculoviral inhibitor of apoptosis repeat-containing 5 (BIRC5), also called survivin, and forkhead box M1 (FOXM1), combined with the CD4^+^ T cell helper peptide universal cancer peptide 2 (UCP2). The peptide mix, that is, the drug product, was emulsified with the adjuvant Montanide ISA 51 VG (Seppic) to obtain a water-in-oil emulsion before subcutaneous administration. Blood collection, standardized PBMCs isolation, and freezing were performed at baseline and then every two or four weeks (**Fig. S5A**). This study was approved by the ethical review boards of all participating institutions and all participants provided informed consent before enrollment in the study.

### Monomer and Tetramer production and validation

Monomers were produced and quality-controlled at the Institute for Immunology, Tübingen, as previously described (12). For tetramer production, monomers were incubated with fluorescently labeled streptavidin -PE (BioLegend), -BV421 (BD), and -BV786 (BD) at 4°C. Following labelling, free streptavidin was blocked by adding biotin at a final concentration of 25µM and incubating on ice for 20min. Finally, the tetramers produced were frozen in a solution of glycerol, NaN_3_, protease inhibitor, and human serum albumin (HSA) diluted in TBS. The tetramers were aliquoted and stored at −80°C until use. No more than 3 rounds of freezing and thawing were performed.

### Tetramer staining assay and memory phenotype analyses in patients

Tetramer staining was performed *ex vivo* on thawed PBMCs and/or cells that had been subjected to *in vitro* stimulation (IVS). Cells were washed with FACS Buffer (PBS supplemented with 2% FCS, 0.02% NaN_3_ and 2mM EDTA) and incubated with two separate tetramer mixes (one containing the OMPs and the other the TAAp-tetramers) each at 2.5µg/ml for 30min at RT. In some experiments, PBMCs were also incubated with three separate mixes, each containing the matched OMP and TAAp tetramers, in order to assess T cell cross-reactivity. The tetramers were diluted in PBS supplemented with 0.02% NaN_3_, 2mM EDTA, and 50% FCS for staining. Afterwards, the cells were washed once with FACS buffer and incubated with mAbs against CD4 (APC-Cy7, BD), CD8 (PE-Cy7, Beckman Coulter), CD14, and CD19 (both PerCP, BioLegend) for 20min at 4°C. A dead live-cell marker (Zombie Aqua, BioLegend) was also included. In the *ex vivo* setting, mAbs against CD45RA (FITC, BD Biosciences) and CCR7 (BV650, BioLegend) were included in the staining mix to evaluate the naïve/memory/effector phenotype of Ag-specific CD8^+^ T cells. After three washes, the samples were analyzed using an LSR Fortessa Cell Analyzer (Becton Dickinson). At least 750,000 and 600,000 cells were acquired in the *ex vivo* and IVS settings, respectively. The data were analyzed using FlowJo v10. Tetramer^+^ T cells were presented as the percentage of cells within living CD8^+^ lymphocytes. For the analysis of the naïve/memory/effector phenotype, the results are shown as the percentage of cells within tetramer^+^ CD8^+^ cells.

### *In vitro* stimulation (IVS) of patient PBMCs

PBMCs from patients were expanded for 12 days (*in vitro* stimulation (IVS)) with the bacterial peptide pool (EO2316, EO2317, and EO2318) before analysis with tetramer staining, IFN-γ ELISpot assay, and ICS (22). For the expansion procedure, PBMCs were thawed in thawing medium (IMDM (Gibco) supplemented with 2.5% heat-inactivated human serum (HS, Capricorn), 1% penicillin/streptomycin solution (PenStrep, Sigma), 50µM beta-mercaptoethanol (β-ME), and 3µg/mL DNase I (Sigma)), washed (1300rpm, 8 min, RT), counted, and resuspended in dedicated medium (IMDM supplemented with 10%HS, 1% PenStrep, and 50µM β-ME, hereafter referred to as TCM). Cells were seeded either in 24 or 48 well plates at approximately 2.5-3.5×10^6^ cells per well and cultured for 24h (37°C, 5%CO_2_). On day 1, peptides were added to the culture medium (final concentration of 1µg/mL for each OMP). On day 3, IL-2 was added to the culture medium (final concentration of 2ng/mL; rhIL-2 reference: R&D 202-IL-010). On day 5, the cells were split by 1/3, and 2ng/mL of IL-2 was again added. On day 7 and 9, media was removed from each well and replaced with fresh TCM containing IL-2 (2ng/mL). If necessary, cells were split 1:2 on day 9. On day 12, the cells were collected, and their viability and number were assessed with an automated cell counter (Nucleo Counter NC-250) using an AO-DAPI staining reagent (solution 18 from Chemometec) (23).

### IFN-**γ** ELISpot assay for patient T cells

For the ELISpot assay, cells after 12 days of IVS with OMPs pools were collected and plated in ELISpot plates (Merck Millipore) pre-coated with a monoclonal anti-IFN-γ mAb (clone 1-D1K purified MabTech) at a density of 200,000 cells/well (24). Cells were either incubated with the human peptide pool (IL-13RA2; BIRC5 and FOXM1 at 5µg/mL each) or the individual bacterial peptide (EO2316 or EO2317 or EO2318, each at 1µg/mL) for 26h. A peptide solvent (10% DMSO in water) was used as the negative control. PHA (10µg/mL) was used as the positive control. After incubation for 26h, cells were discarded and detection of IFN-γ was performed by adding the anti-human IFN-γ biotinylated mAb (clone 7-B6-1, MabTech) for 2h, followed by incubation with ExtrAvidin Alkaline Phosphatase (Sigma) for 1h, and finally adding the substrate BCIP/NBT (Sigma) to the wells. Plates were left to dry for at least 12h at RT in the dark prior to plate imaging and spot counting using an Immunospot® S6 UNIVERSAL Analyzer (C.T.L). The number of spots in peptide-stimulated wells (n=3 for each condition) was compared to that obtained in the negative control (solvent control wells) using a permutation test (distribution-free resampling DFR2x) (25). If less than three replicates per condition were available, analyses were performed manually based on positivity criteria from the laboratory (at least twice the spot count in peptide-stimulated wells compared to solvent control plus a minimum of 6 spots per 100,000 cells seeded).

### Intracellular staining assay (ICS) in patient PBMCs

1.5-2×10^6^ cells collected after 12 day-IVS were seeded in a round-bottom 96-well plate for assessment of the cytokine profile. For ICS, cells were stimulated for 1h in TCM with peptides (EO2316, EO2317, EO2318, IL13RA2, BIRC5, and FOXM1 at 10µg/mL each) along with anti-CD107a (BD Biosciences, FITC, clone H4A3) to capture degranulation. After 1h, 10μg/mL brefeldin A (Sigma-Aldrich) and GolgiSTOP (BD Biosciences) at a dilution of 1:1500 were added to each well, and the cells were incubated at 37°C for 14 h. After stimulation, cells were washed and stained for 20 min at 4°C with mAbs CD4-APC-Cy7 (BD, clone RPA-T4), CD8-BV605 (clone RPA-T8), CD14-PerCP (Clone 63D3) and CD19-PercP (clone HIB19) all BioLegend diluted in FACS buffer (PBS + 2% heat-inactivated (hi) fetal bovine serum (FBS) + 0.02% NaN3 + 2 mM EDTA). In addition, a live cell dye (Zombie Aqua, Biolegend) was added to discriminate the dead cells. After washing with FACS buffer, the cells were fixed and permeabilized with BD Cytofix/Cytoperm (BD Biosciences) for 20 min at 4°C, followed by staining for the intracellular markers IFN-γ (Human anti-IFN-γ, AF700, clone B27), TNF (Human anti-TNF, PaBl, clone Mab11), IL-2 (anti-human IL-2, PE-Cy7, clone MQ1-17H12), and CD154 (anti-human CD154, APC, clone 24-31) all from Biolegend in Perm wash buffer (PBS + 0.02% NaN_3_ + 0.5% bovine serum albumin (BSA) + 0.1% saponin) for 20 min at 4°C. After three washes, the samples were analyzed using a BD LSRFortessa Cell Analyzer (BD). At least 4×10^5^ cells were collected. The data were analyzed using FlowJow v10. Results are presented as the percentage of cells within the CD8+/CD4-cells. Polyfunctionality analysis was performed using Pestle Version 2.0 and Spice Version 6.1 (26).

### Statistical Analysis

Results are expressed as mean ± SEM or SD. The Mann-Whitney U test was used to compare the two groups. Comparisons between tumor growth curves were performed using a two-way ANOVA test, and multiple comparisons were corrected using the Bonferroni coefficient. Statistical significance was determined using Prism software (GraphPad Software). Statistical significance was set at P < 0.05.

### Materials, Data and Resource availability

Requests for additional information, resources, reagents and unique materials developed in this study should be directed to the corresponding author and subject to a material transfer agreement. All relevant data are available in the main text and the supplementary materials.

## Results

### A Comprehensive Multistep Process for Discovering and Selecting OMPs for Cancer Immunotherapy

We developed a comprehensive bioinformatics pipeline to discover commensal-derived peptides (CDPs) that exhibit molecular mimicry (sequence similarities) with tumor-associated antigen peptides (TAAps). This approach targets peptides with enhanced affinity for HLA class I molecules, particularly HLA-A2, due to its prevalence in approximately 49% of the Caucasian population, with a significant majority expressing the HLA-A*02:01 allelic variant (27). Our strategy aims to identify immunogenic CDPs capable of initiating cross-reactive CD8+ T cell responses against cancer cells presenting these TAAps. We screened 224 HLA-A2 verified cancer antigen peptides derived from 113 well describe TAAs against a large public database of gut commensal proteins (**Supplementary Table S1 and see M&M**). This database comprises almost 10 million genes from 1267 individuals, highlighting the potential of the gut microbiome as a source of short peptides that can potentially mimic all described MHC class I tumor peptides (28). Peptide pairs (CDP/TAAp) were selected according to criteria aimed at maximizing amino acid (aa) changes that improve HLA-A2 binding affinity (anchor positions), while maintaining strict identity within the central aa sequence (core positions) crucial for TCR recognition when presented by MHC class I (**Fig. 1A**) (29).

**Figure 1.**
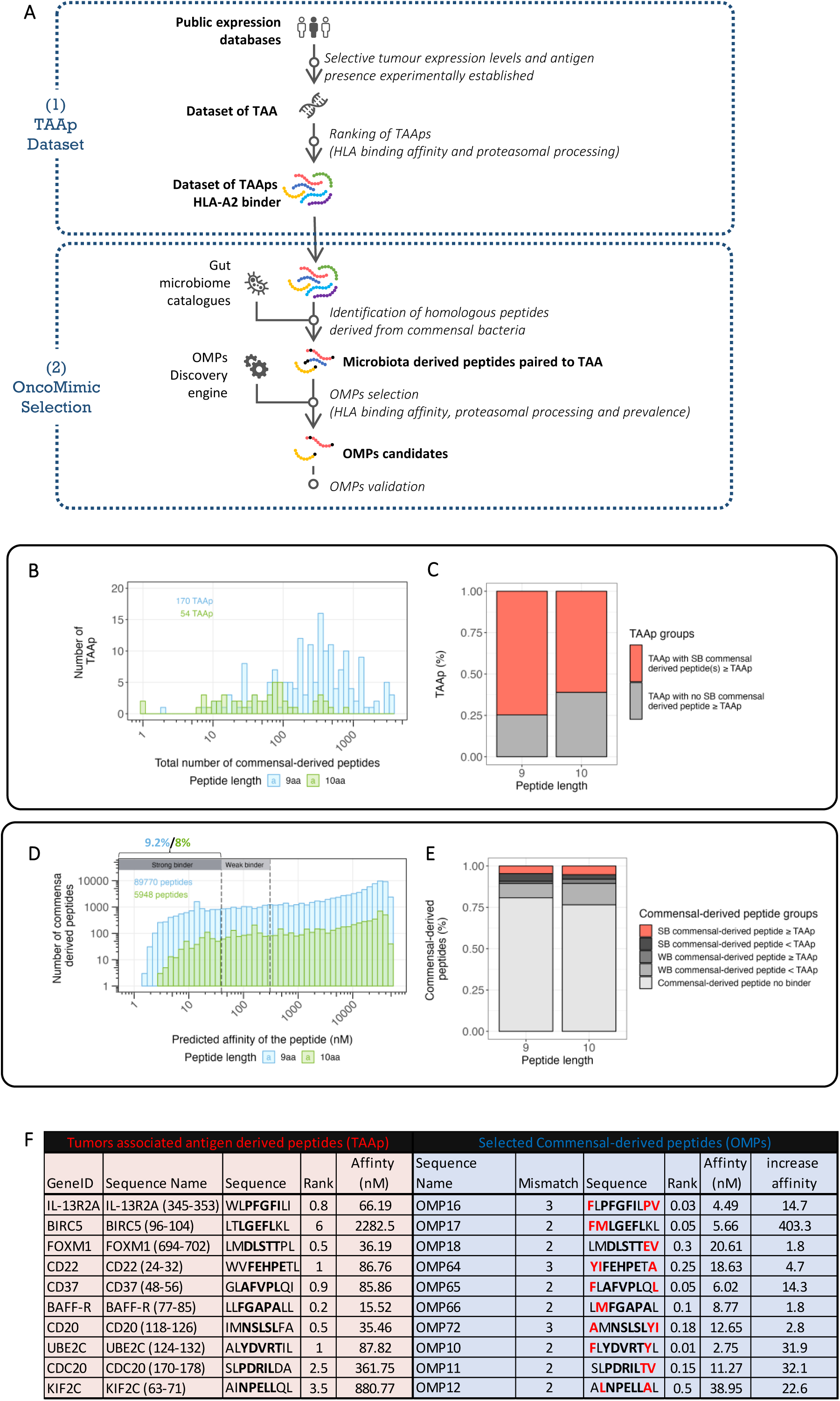
Discovery and selection process of OMPs for cancer immunotherapy. **<u>A</u>**<u>. OMPs discovery: A two-step process.</u> (1) A dataset of HLA-A2 peptides derived from tumor-associated antigens (TAAs) was generated from human public databases according to several parameters. (2) OMP candidates were identified by scanning each TAA-derived peptide against all peptide sequences of the same length from gut microbiome catalogues using different homology criteria (detailed in M&M). **<u>B</u>**<u>. TAA derived peptides distribution based on the number of their commensal-derived peptides:</u> The graph shows the distribution of 224 TAA-derived peptides (TAAps) based on the number of their associated CDPs. **<u>C</u>**<u>. Repartition of TAA-derived peptides based on the HLA-A2 affinities of their commensal-derived peptides:</u> The graph represents the repartition of TAA-derived peptides based on two categories: TAAp with strong binder (SB) CDPs ≥ TAAp or TAAp with no SB strong binder CDPs ≥ TAAp. In other words, the presence (in red) or absence (in grey) for each TAAp of at least one strong binder CDP, stronger than their TAAp counterparts. In the legends of Figures 1C and 1E, the symbol ‘≥’ indicates ‘a stronger predicted affinity than,’ while ‘<’ signifies ‘a weaker predicted affinity than.’ **<u>D</u>**<u>. Commensal-derived peptide distribution based on HLA-A2 predicted binding affinities:</u> The graph shows the distribution of commensal-derived peptides based on their HLA-A2 predicted binding affinity. Commensal-derived peptides with the highest affinities (lower nM values) are represented on the left side of the graph. We classified those CDPs in 3 categories based on their %rank (according NetMHCpan3.0 recommendations): Strong binders (SB) are defined as having %rank<0.5, weak binders (WB) with %rank<2 and no binder peptides with %rank>2. **<u>E</u>**<u>. Commensal-derived peptide repartition based on their HLA-A2 predicted binding affinity compared to their TAAp counterparts:</u> The graph shows how commensal-derived peptides are categorized into five groups: SB commensal-derived peptide with an HLA-A2 affinity equal to or stronger than the associated TAAp; SB commensal-derived peptide with an HLA-A2 affinity equal to or weaker than the associated TAAp; WB commensal-derived peptide with an HLA-A2 affinity equal to or stronger than the associated TAAp; WB commensal-derived peptide with an HLA-A2 affinity equal to or weaker than the associated TAAp; Commensal-derived peptides that are no binders to HLA-A2. **<u>F.</u>** <u>Characteristics of the ten selected HLA-A2 OMPs in this study and their homologous TAAps: Gene ID of the targeted TAA, sequence name</u> of TAAp and corresponding OMP. Amino acid (aa) sequences predicted affinity rank and binding affinity (nM) to HLA-A*02:01 molecule of each pair and the number of mismatches are shown. Amino acid substitutions are indicated in red and in dark bold are the 5 aa of the peptide core. Affinity fold increases between OMP and its corresponding TAAp.

Our screening identified 95,718 CDPs as potential matches, with the number of CDPs per TAAp ranging from 1 to 3,543 (**Supplementary Table S2 & Fig. 1B**). 74.7% of the 9-mers and 61.1% of the 10-mers from the selected TAAps had at least one strong binder (SB) HLA-A2 CDP with enhanced binding properties demonstrating that a majority of the well-documented HLA-A2 TAAp have at least 1 strong binder CDP equivalent (**Fig. 1C**). Among all CDPs, 9.2% of the 9-mers and 8.0% of the 10-mers were predicted to be SB (**Fig. 1D**), with 4.6% predicted to have higher affinities than their TAAp counterparts. We designated this high-affinity CDPs subset as OncoMimic^™^ peptides (OMPs) meeting our criteria for sequence homology with TAAps and exhibiting increased HLA-A2 affinities (**Fig. 1E**).

Ten OMPs were selected to assess their biological potential. Six OMPs-matched TAAps derived from overexpressed tumor antigen (IL-13RA2, BIRC5, FOXM1, UBE2C, CDC20 and KIF2C) prevalent in solid tumors like glioblastoma, lung, breast, and colorectal cancers and identified as key tumor drivers (30–35). The remaining four OMPs-matched TAAps derived from CD22, CD37, BAFF-R, and CD20, B cell surface markers described as targets in B cell lymphoma therapies (36). These ten OMPs were selected from a comprehensive pool of candidates, distinguished as the most promising based on several parameters. This included sequence similarity to TAAps, ensuring minimal mismatches to maintain molecular mimicry; high HLA-A2 binding affinity, predicting potent immunogenicity; effective proteasomal cleavage potential, suggesting their likely processing and presentation at the gut level; and prevalence within the gut microbiome, suggesting their influence on the human T-cell repertoire (**Supplementary Table S2 and M&M for details**). These carefully selected OMPs present 2 or 3 mismatches and exhibit significant enhanced predictive affinities ranging from 1.8 to 400-fold higher than their TAAp counterparts (**Fig. 1F**). This selection process aims to predict that these OMPs can trigger strong immune responses in a wide population segment.

### OMPs induce cross-reactive T cell responses *in vivo*

To validate the *in silico* HLA-A2 affinity predictions, we used TAP-deficient, HLA-A*02:01-positive T2 cell line. Those cells provided an established model to assess peptide-HLA-A2 binding measured by minimizing endogenous peptide competition (**Fig. S1A**). Our *in vitro* binding assays revealed that each tested OMP exhibited enhanced binding affinity compared to their corresponding TAAp (**Fig. 2A, S1B and S1C)**. Stability assays further highlighted that OMP/HLA-A2 complexes were significantly more stable than the TAAp/HLA-A2 counterparts, with dissociation half-life complex (DC50) values ranging from 2h to more than 24h. (**Fig. 2B, S1B and S1D**). This enhanced stability suggests a prolonged antigen presentation, potentially improving T-cell recognition and response. The immunogenicity capacity as well as the ability of the OMPs to induce cross-reactive T cells were subsequently investigated *in vivo* using humanized HLA-A2-DR1 transgenic mice. Upon OMP immunization, antigen-specific IFN-γ T cell responses were evaluated (**Fig. 2C**). Robust T cell responses were observed for all OMPs tested, while no T cell activation was detected in naïve mice. The induction of cross-reactive T cell responses were further confirmed by the reactivity of those OMP-induced T cell to produce IFN-γ upon *ex vivo* restimulation with the corresponding TAAps, underlining the capacity of those OMPs in eliciting OMP-/TAAp-specific T cells (**Fig. 2D and Fig. S1E**). The numbers of TAAp- and OMP-responding T cells in individual mice strongly correlated, with a Pearson coefficient ranging from 0.42 for the least correlated peptide pair, namely OMP10/UBE2C, to 0.99 for the most correlated pairs (7 out of 10 peptide pairs) **(Fig. S1F**). Interestingly, we noted differences in cross-reactivity levels between the 10 OMP/TAAp pairs varying from 11% to 17% for the least cross-reactive pairs (OMP16/IL-13RA2 and OMP18/FOXM1, respectively) to 95% for the most cross-reactive pair (OMP17/BIRC5). Overall, these results revealed the capability of OMPs to induce strong IFN-γ-producing T cells responses recognizing both OMP and TAAp.

**Figure 2.**
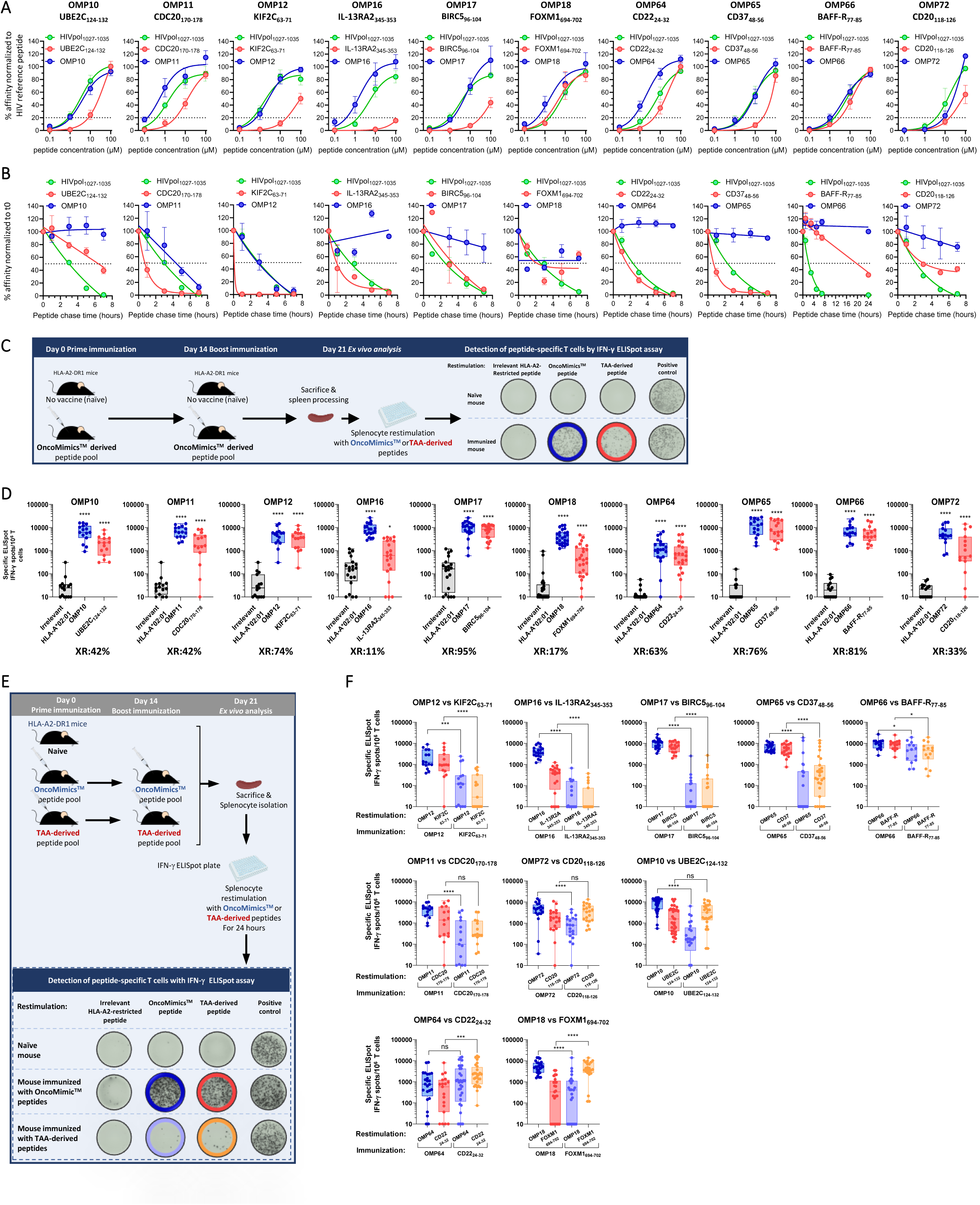
OMPs selected *in vitro* for their HLA-A2 binding and stability properties and *in vivo* for their immunogenicity are capable of eliciting potent cross-reactive responses in HLA-A2/DR1 mice. **<u>A.B</u>**<u>. Binding and stability of TAA-derived peptides and their OMP counterparts.</u> Comparison of HLA-A2 binding (A) and stability (B) of TAA-derived peptides (red), their OMP counterpart (blue) and HIV reference peptide (green) in the T2-binding assay. Data shown represent four to seven independent experiments. Symbols represent the mean and error bars indicate SEM. **<u>C.</u>** <u>Schematic representation</u> of the *in vivo* experimental set-up used to assess OMPs immunogenicity and CD8^+^ T cell-dependent cross-reactive response against TAAps. **<u>D.</u>** *<u>In vivo</u>* <u>immunogenicity and cross-reactivity.</u> The frequency of peptide-specific T cells that produce IFN-γ was determined using ELISpot analysis of splenocytes from mice vaccinated with the indicated peptides. In grey, blue and red are shown the negative control, OMP and TAAp, respectively. Below each graph, the percentages of cross-reactivity (XR) between the OMP and its TAA counterpart are displayed. **<u>E.</u>** <u>Schematic representation</u> of the *in vivo* experimental set-up used to compare the capacity of OMPs and TAA-derived peptides to induce a TAA-specific response. **F.** <u>OMP- and TAAp-induced cross-reactive responses.</u> Comparison of OMP efficacy in inducing a TAA-specific response to that of their TAA counterparts in A2/DR1 humanized mice, assessed on splenocytes from mice immunized with the indicated peptides by IFN-γ ELISpot assay. In dark blue, red, light blue and orange are shown the OMP vaccination and restimulation condition, OMP vaccination and TAAp restimulation, the TAAp vaccination and OMP restimulation condition and the TAAp vaccination and restimulation condition, respectively. The data shown in **D.** and **F.** are from one to five independent experiments, symbols indicate individual mice (n= 5-25 mice) and bars represent min and max values. Statistical comparison was performed using an unpaired non-parametric test (Mann-Whitney). * p < 0.05, ** p < 0.001, **** p < 0.0001, ns: non-significant.

Mice were then immunized with either the OMP or the TAAp counterpart, and T cells were restimulated *ex vivo* with either of the 2 peptides (**Fig. 2E**). T cell responses against KIF2C, IL-13RA2, BIRC5, CD37, and BAFF-R peptides were weaker when the mice were immunized with TAAp compared to their OMP counterparts. This demonstrates better immunogenicity for OMP12, OMP16, OMP17, OMP65, and OMP66 over their respective homologous TAAps (**Fig. 2F**, top panel). CDC20-, CD20- and UBE2C-specific T cell responses were of similar magnitude, irrespectively of the initial vaccination (**Fig. 2F**, middle panel). Only CD22- and FOXM1-specific T-cell responses were lower after immunization with OMPs than after immunization with TAAps (**Fig. 2F**, bottom panel).

### OMPs trigger CTL responses with antitumor activity

The *in vivo* cytotoxicity of the OMP-specific T cells was investigated using a fluorescence-based cytotoxic T-cell assay (37). Following a prime/boost immunization of A2/DR1 mice with the OMP pools (OMP17, OMP18, OMP10, OMP11, OMP12) or (OMP64, OMP65, OMP66, OMP72), a 1:1 mix of syngeneic splenocytes loaded with the corresponding OMPs (bright labelling) and unloaded (dim labelling) were injected into immunized or control animals (**Fig. 3A**). Flow cytometry, conducted 20h post-injection, demonstrated that OMP-specific T cells exhibited substantial cytotoxic activity towards target cells without affecting control cells viability indicating that OMPs elicit targeted and robust CTL responses (**Fig. 3B**). CTL activity was observed against T2 cells loaded with most of the individual OMPs tested, although the responses against the OMP64 were lower and those to OMP72 were negligible (**Fig. S2A and S2B**). More importantly, these OMPs triggering CTL activities were efficacious against splenocytes pulsed with the matched TAAp pool (BIRC5, FOXM1, UBE2C, CDC20, and KIF2C) or (CD22, CD37, BAFF-R, and CD20) (**Fig. 3C**) proving their cross-reactivity.

**Figure 3.**
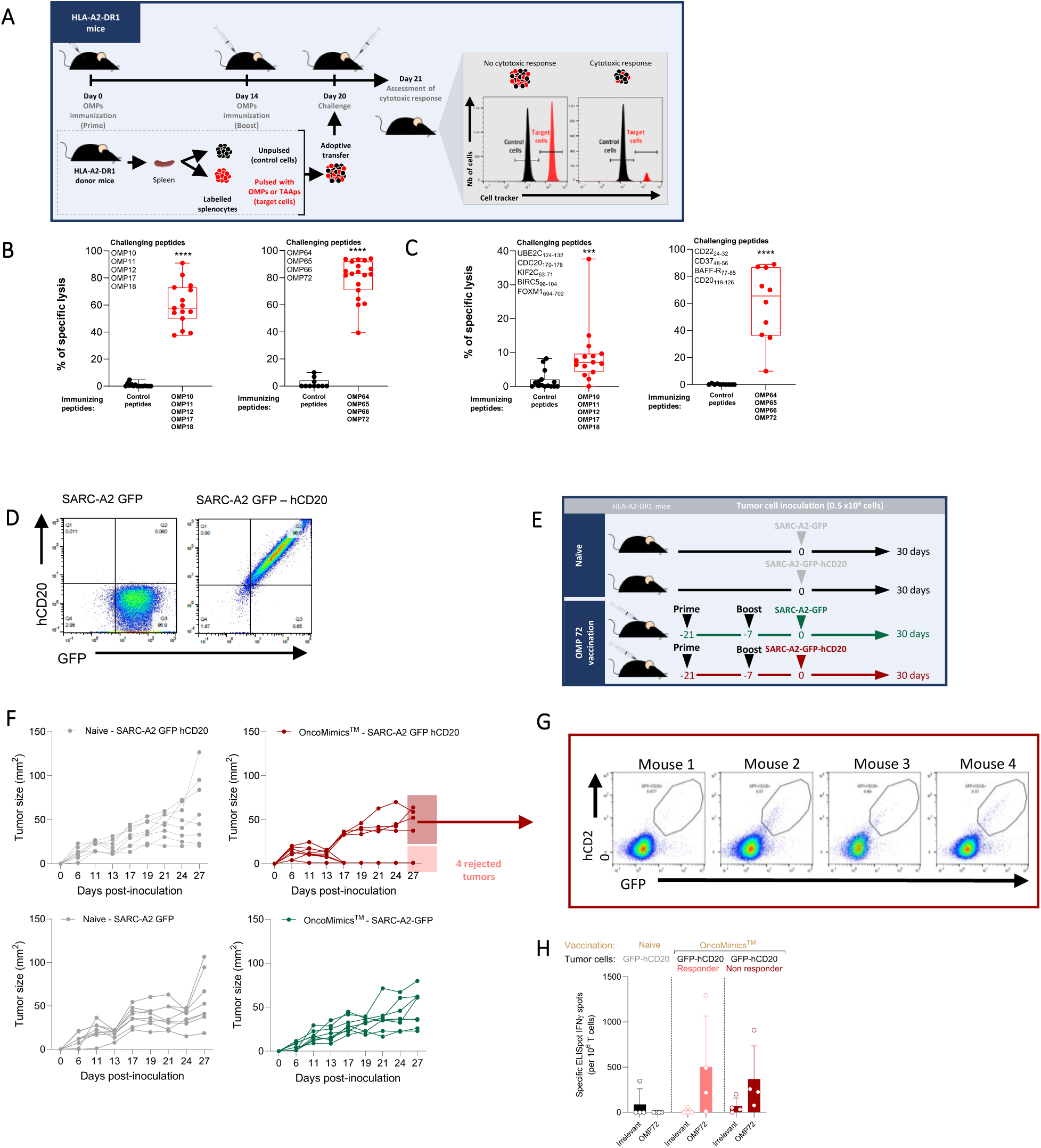
OMPs-based vaccines elicit functional cytotoxic T cells in mice that are cross-reactive against human TAA-derived peptides *in vivo*. **<u>A.</u>** <u>Schematic representation of the experimental setup employed to assess T cell cytotoxicity elicited *in vivo* by OMPs.</u> HLA-A2-DR1 mice were vaccinated with OMPs and challenged post-vaccination with syngeneic splenocytes labeled with cell tracking dye. Target T cells (red) were pulsed with a mix of OMPs or TAA-derived peptides and mixed at equal ratio with unpulsed control cells (black) before being adoptively transferred into immunized mice. **<u>B.C.</u>** *<u>In vivo</u>* <u>cytolytic activity of OMP-induced T cells against OMP- or TAAp-pulsed targeT cells.</u> Percentages of *in vivo* specific lysis of splenocytes pulsed with the indicated pool of OMPs (**B**) or TAA-derived peptides counterparts (**C**) (challenging peptides) after immunization with the indicated peptide pool or control peptides (immunizing peptides). Data shown in **B** and **C** are from two to four independent experiments, symbols indicate individual mice (n= 10-20 mice) and bars represent min and max values. Statistical comparison was performed using an unpaired non-parametric test (Mann-Whitney). *** p < 0.001, **** p < 0.0001. **<u>D.</u>** <u>Flow cytometry analysis performed on SARC-A2</u> GFP (left panel) and on SARC-A2-GFP-CD20 (right panel) sarcoma cells showing post-transduction expressions of GFP and hCD20 proteins. **<u>E.</u>** <u>Schematic representation of the experimental setup used to evaluate the antitumor effect of OMPs.</u> Mice were immunized with OMP72 using a prime-boost administration regimen. 21 days post-prime immunization, mice were inoculated with hCD20-GFP or GFP-expressing SARC-A2 sarcoma cells and tumor growth was monitored. **<u>F.</u>** <u>Tumor kinetics on individual mice over time for each group</u>. Tumor size measurement (mm^2^) in A2/DR1 naïve or OMP vaccinated mice engrafted with 0.5×10^6^ SARC-A2-hCD20-GFP or SARC-A2-GFP tumor cells (n=8 mice) is shown oven 30 days. **G.** h<u>CD20 expression assessment in vaccinated group</u>. Flow cytometry dot plots showing the expression of human CD20 and GFP in SARC-A2 cells extracted from whole tumors at day 30 on animals still bearing tumors (n=4). **H.** <u>Vaccine-specific induced T cell responses.</u> The frequency of peptide-specific T cells producing IFN-γ per million splenic T cells was determined by ELISpot at day 30.

To further explore the *in vivo* activity of OMP-specific T cells against tumors, we established a tumor regression model using the syngeneic HLA-A2^+^ sarcoma cell line (SARC-A2), engineered to express the human CD20 antigen as well as the GFP (SARC-A2-GFP-hCD20) or the GFP alone control cell line (SARC-A2-GFP) (**Fig. 3D**). This approach allowed to assess OMP72 capacity to trigger CD20 targeted CTL-mediated tumor cell killing. Post-immunization with OMP72, A2/DR1 mice were engrafted with the SARC-A2 tumor cells and exhibited a significant reduction in tumor volume compared to control groups, with half of the SARC-A2-GFP-hCD20 mice group achieving complete tumor regression (**Fig. 3E and 3F**). Interestingly, we observed that mice for whom tumor control was not achieved, there was a loss of CD20 surface expression on tumor cells, suggesting a potential escape mechanism from OMP72-specific T cell surveillance (**Fig. 3G**). Supporting this hypothesis, comparable levels of OMP72-specific IFN-γ-producing cells were measured across both responsive and non-responsive mice, indicating the immunogenicity of the OMP72 peptide (**Fig. 3H**). This successful tumor regression exemplifies the capability of OMPs to foster potent and targeted anti-tumor CTL responses.

### OMP-specific T cells represent a prevalent pool of T cells in the human population

The capacity of the selected OMPs to induce efficient TAA-specific cross-reactive CTL responses was then evaluated in humans peripheral blood mononuclear cells (PBMCs) isolated from HLA-A2^+^ healthy donors (HDs) and stimulated *in vitro* in the presence of OMPs (**Fig. 4A**). Flow cytometry analysis using peptide-MHC tetramers revealed that all the OMPs induced cross-reactive OMP-/TAAp-specific CD8^+^ T cells (**Fig. S3A, S3B and S3C**). Within the same OMP/TAAp pairs, the extent of cross-reactivity varied between different HD PBMCs ranged from 10% to 100% of cross-reactivity. In addition, the degree of cross-reactivity among OMP/TAAp pairs showed variability across the different pairs. Specifically, some OMP/TAAp pairs showed cross-reactivity levels below 33% (OMP10, OMP11, OMP18, OMP64), some between 33 and 66% (OMP16, OMP65, OMP66, OMP72), and some above 66% (OMP12, OMP17), in average when compared to the total elicited OMP-specific T cell responses (**Fig. S3D**). This pattern emphasizes the variability not only within individual responses to the same OMP/TAAp pair but also across different OMP/TAAp pairs, suggesting that multiple factors influence the properties of individual OMP immunogenicity.

**Figure 4.**
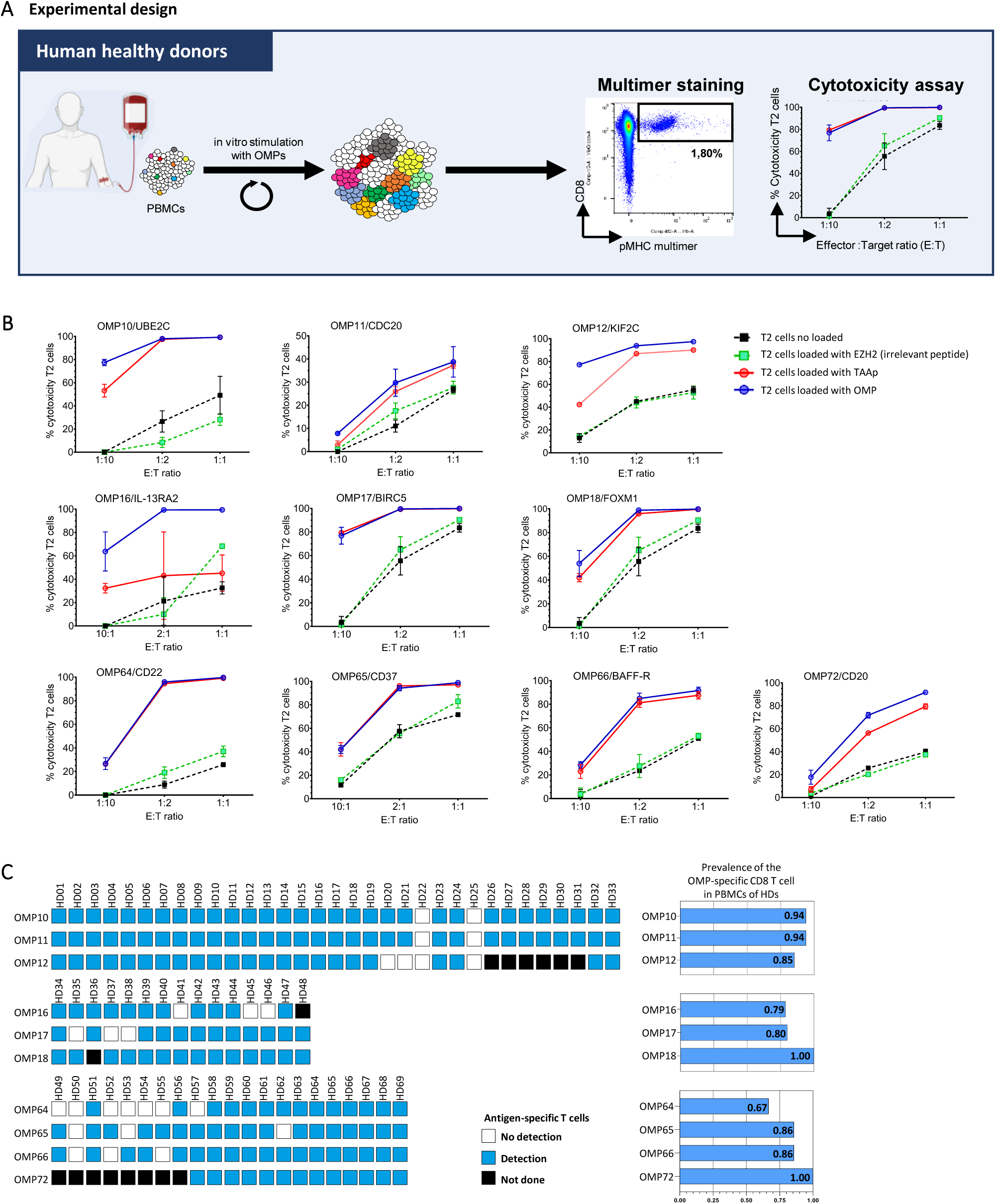
OMP-specific human T cells recognize TAAs and exert specific cytolytic activity. **A.** Schematic overview for determining antigen-specific CD8^+^ T cell response using healthy volunteer PBMCs and *in vitro* expansion of CD8^+^ T cells as described in *Material and Methods*. **B.** Cytolytic activity of the OMP-specific human CD8^+^ T cell. CTL killing activity assessed against T2 cells loaded with the OMP and the TAA-derived peptide (TAAp) counterpart after 24 h incubation. Each graph displays representative data from individual healthy donors. Cytotoxicity percentage (y-axis) and effector:target ratio (E:T ratio) (x-axis) are indicated on the graph. EZH2-B2 irrelevant peptide and unpulsed T2 cells are also shown as negative controls. Error bars represent mean +/- standard deviation (SD). The solid lines depict T2 cells loaded with bacterial peptides (OMPs in blue) and human derived peptides (TAAp in red) and the dashed lines represent controls (unloaded in black, EZH2-loaded in green). Cytotoxicity percentage was calculated as specified in *Materials and Methods*. **<u>C.</u>** Prevalence of OMP-specific CD8^+^ T cells in healthy donors (HD). Prevalence of OMP-specific CD8^+^ T cells in HDs was determined by surface staining using pMHC tetramers after expansion of PBMCs using OMPs and is shown on histogram graph. Left panel shows individual detection of OMPs in each HD with white, blue and black squares representing HDs in which OMPs were not detected, detected or no analyses was performed, respectively. Right panel shows the general frequency of OMP detection in the human healthy population. The number of analyzed HDs is indicated into brackets for each OMP: OMP10 and OMP11 (n=33), OMP12 (n=27), OMP16 (n=14), OMP17 (n=15), OMP18 (n=14), OMP64, OMP65, OMP66 (n=21) and OMP72 (n=13).

The functional cytotoxic capability of OMP-stimulated PBMCs was provided by assessing their killing activity on T2 cells loaded with OMPs. We observed cytotoxic activity across all OMPs, confirming the functional efficacy of OMP-specific T cell in recognizing and destroying target cells that present these peptides. This specificity was confirmed as control T2 cells, either pulsed with an irrelevant peptide or left unloaded, showed negligible or significantly reduced killing. Additionally, the cross-reactive cytotoxic capability of the OMP-stimulated PBMCs was evident, as they also targeted T2 cells presenting matched TAAps. Reduced and variable cytotoxic values were observed with IL-13RA2, likely attributable to the low affinity of IL-13RA2 peptide and poor stability when artificially presented by T2 cells (**Fig. 4B and S1B**). In most cases, killing occurred at a low effector to target cell (E:T) ratio of 1:10, strengthening the efficacious functional cytotoxic activity of these effector cells. Moreover, OMP-specific T cells killed OMP- and TAAp-loaded T2 cells with equal efficiency (**Fig. 4B**). We further determined the prevalence of OMP-specific CD8^+^ T cells in the human population. Notably, all OMP-specific CD8^+^ T cells were detected in over 79% of HLA-A2^+^ PBMCs samples, except for OMP64 which showed a prevalence of 67% (**Fig. 4C**). This high detection rate emphasizes the widespread potential of OMPs to engage the immune system across a broad segment of the population, highlighting their relevance for immunotherapeutic strategies against cancer.

### OMPs induce fast, strong and long-lasting polyfunctional T cell responses in cancer patients with recurrent glioblastoma

Immune responses induced by OMPs were evaluated in patients from the EOGBM1-18/ROSALIE clinical trial (NCT04116658). EOGBM1-18 is a first-in-human, phase Ib/IIa trial in patients at first recurrence of glioblastoma after a radiotherapy/temozolomide regimen. Patients received an immunotherapy, designated EO2401, comprising three OMPs (EO2316/OMP16, EO2317/OMP17, and EO2318/OMP18) and the universal cancer peptide 2 (UCP2), administered in combination with nivolumab (**Fig. S4A**). Here, we present data from the first three patients included in Cohort 1 [multiple doses of EO2401 monotherapy followed by continued EO2401 in combination with nivolumab], designed to evaluate the safety and the tolerability of the approach. Immune responses were evaluated in cryopreserved PBMCs both *ex vivo* and following *in vitro* stimulation (IVS). We performed peptide/MHC tetramer staining, intracellular cytokine staining (ICS), and IFN-γ ELISpot assays.

We initially evaluated the functional capacity and cross-reactivity of antigen-specific memory and effector T cells induced by EO2401 treatment using short-term IVS with OMPs, followed by MHC tetramer staining. This experimental setup effectively recalls memory and effector cells, enabling detailed analysis of their responses (38,39). Tetramer staining revealed the presence of specific T cells for the three OMPs with various frequencies among patients. EO2317 and EO2318 were the most immunogenic peptides, with significant T cell responses in all three patients. After *in vitro* stimulation of PBMCs with OMPs and a short expansion, we observed at the plateau of the response (6 weeks) a marked proliferative capacity of OMP-specific T cells with approximately 80%, 35%, and 28% of total CD8^+^ T cells for Patients #1, #2, and #3, respectively (**Fig. 5A**, gating strategy and representative staining in **Fig. S4B-C**). Long-term follow-up of Patients #1 and #2 revealed that the strong immune response was sustained until week 44 (**Fig. 5A**). Interestingly, in Patients #2 and #3, we identified pre-existing OMP-specific CD8^+^ T cells that increased more than 60-fold upon treatment (*ex vivo* vs. post-IVS tetramer staining) (**Fig. 5B**). BIRC5- and FOXM1-specific CD8^+^ T cells were readily detected in IVS with OMPs. In contrast, frequencies of cross-reactive T cells against IL-13RA2 were low in Patient #1 and below the detection limit in Patients #2 and #3 (**Fig. 5C** and representative staining in **Fig. S5D)**. At the plateau, we observed 40% and 14% of BIRC5-specific CD8^+^ T cells and approximately 1% of FOXM1-specific CD8+ T cells for Patients #1 and #2, respectively. The long-lasting and strong immune responses against BIRC5 and FOXM1 followed the same kinetics as EO2317- and EO2318-specific responses, respectively (**Fig. 5A** and **5C**). No pre-existing TAA-specific cells were detected, except for Patient #2, where pre-existing BIRC5 reactivity was observed (**Fig. 5D and S4D**). We then assessed the OMP-/TAA-specific T cell cross-reactivity by performing tetramer staining on post-IVS PBMCs using the tetramer pairs EO2317/BIRC5 and EO2318/FOXM1 simultaneously. Remarkably, all TAA-specific CD8^+^ T cells in the tested patients (Patients #1 and #3) also bound the matched OMP tetramer T cells (**Fig. S4E**). This result strongly suggests that the OMP-specific T-cell population serves as precursor for the generation of TAA-specific T cells. Taken together, we conclude that peptide vaccination with the OMPs EO23016, EO2317, and EO2318 can efficiently mount a rapid, strong OMP-specific T cell response, with high proliferative T cells upon OMP re-exposure, and ability to cross-react with the targeted TAAps (BIRC5, FOXM1, and to a lesser extent IL-13RA2). Corroborating these findings, only OMP-stimulated PBMCs from Patient#1 resulted in the proliferation of TAAp-specific T cells, unlike stimulation with the TAAp. Further, neither increasing the concentration of TAAps nor adding cytokines known to support antigen-specific memory T cell (IL-7, IL-15 and IL-21) do not lead to the proliferation of TAAp-specific T cells, underscoring OMP superior efficacy in driving memory T cell proliferation (**Fig. S4F and S4G**). Next, we performed functional characterization of these OMP-/TAAp-specific cross-reactive CD8^+^ T cells. We used two functional assays (IFN-γ ELISpot and ICS) following IVS with OMPs. High levels of IFN-γ T cell responses against the three OMPs and the TAAp pool were detected in Patients #1 and #2 and, to a lesser extent, in Patient #3. (**Fig. 5E and 5F**). As expected from MHC tetramer staining performed after IVS (**Fig. 5A**), we also observed pre-existing reactivity against EO2316 in Patient #3 (**Fig. 5E**). The functionality of antigen-specific CD8^+^ T cells was confirmed. The production of Tc1 cytokines (IFN-γ and TNF-α), a Lamp1/CD107a CTL marker, and cell activation/proliferation markers (IL-2 and CD154) were evaluated. OMP- and TAA-polyfunctional IFN-γ^+^-, TNF-α^+^-, IL-2^+^-, CD154^+^-, and CD107a^+^-specific CD8^+^ T-cell responses were observed. In line with the tetramer staining results, T cell polyfunctionality was mainly observed upon restimulation with EO2316, EO2317, EO2318, BIRC5, and to a lesser extent, with FOXM1 (**Fig. 5G and** representative staining in **Fig. S4H**).

**Figure 5.**
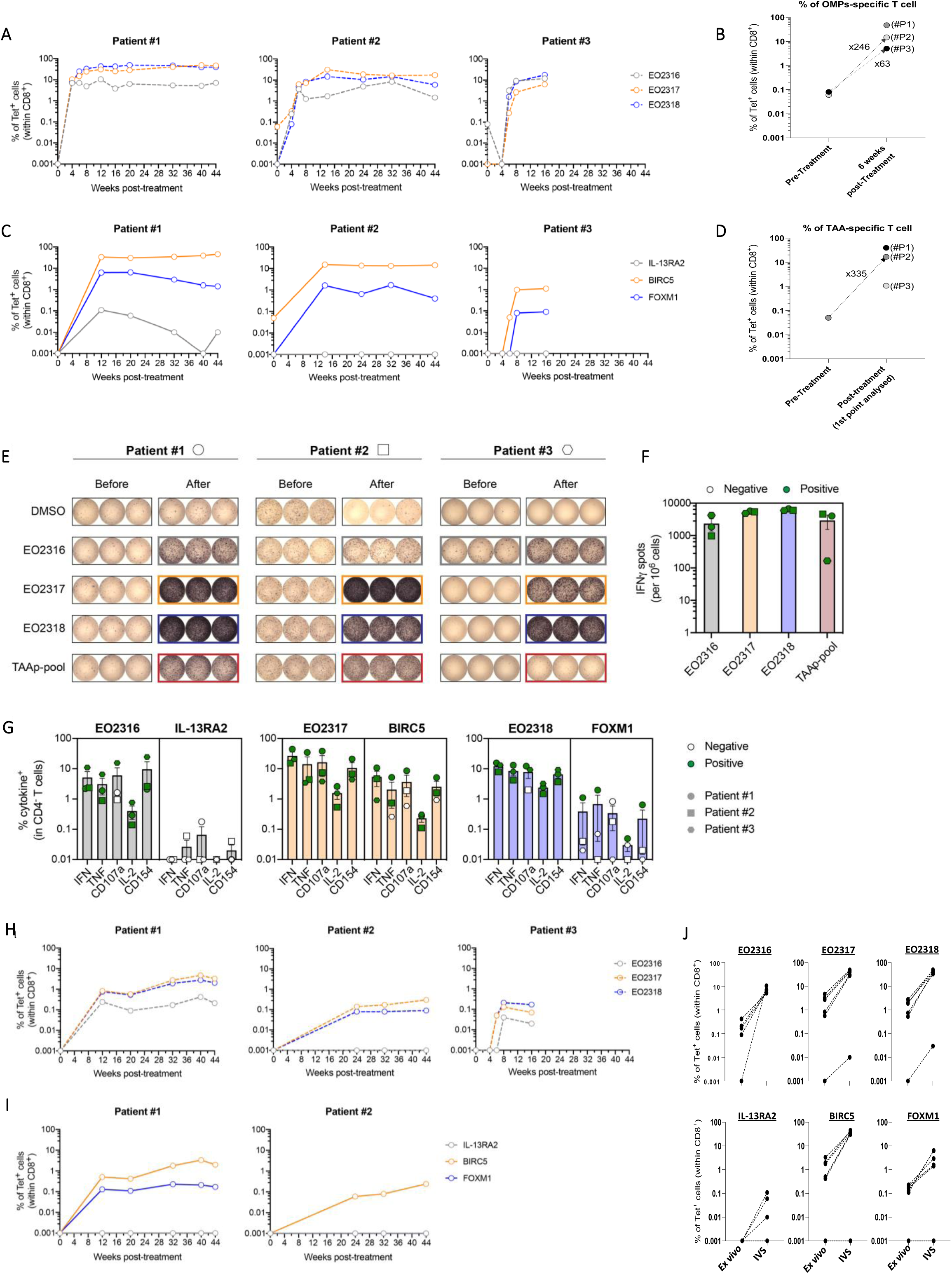
Induction of OMP-specific and TAA-cross-reactive T cell responses in glioblastoma patients post-vaccination. <u>Post-IVS T cell responses to OMPs or TAAps from patients are shown in A-D.</u> **A**, **C**: OMP-(**A**) and TAA-(**C**) specific CD8^+^ cells detected using EO2316-, EO2317-, EO2318-, IL-13RA2-, BIRC5-, and FOXM1-tetramers on PBMC samples subjected to 12 days *in vitro* stimulation (IVS) with the OMP peptide pool. Data obtained from all available samples of patients from cohort 1 (n=3), each dot represents the percentage of tetramer-positive CD8^+^ cells for the respective patient at the defined week. Negative values are plotted as 0.001. **B, D**: <u>Immune responses from patients before and after treatment</u>. Total OMP-(**B**) and TAAp-(**D**) specific CD8^+^ T cells detected in patient PBMC samples before and after treatment at the indicated time points. Data represent the added percentages (%) of all OMP (**B**) or TAAp (**D**), determined by multimer staining shown in **A** and **C**. A fold increase was calculated and is shown in the graph when OMPs- or TAA-specific T cells were detected before treatment. **E-F.** <u>Generation of functional antigen-specific cells after vaccination</u>. Immune responses of patients before and after treatment were analyzed using IFN-γ ELISpot post-IVS. IFN-γ ELISpot wells are shown for all three patients before and after vaccination, with positive wells highlighted with colored squares (**E**). Quantification of the number of spots after vaccination for each patient, as well as the mean ± SEM, are shown in (**F**). The IFN-γ spots were normalized to 10^6^ cells after background subtraction (negative control, DMSO). **G**. <u>Polyfunctional T cells are generated by OMP vaccination.</u> ICS quantification of the % of activation marker-(IFN-γ, TNF, CD107a, IL-2, and CD154)-positive T cells upon stimulation with the indicated peptides (top of each bar graph). For **F** and **G**, Patients #1, #2, and #3 are shown as circles, squares, and hexagons, respectively. The symbols filled in green and white represent positive and negative responses, respectively (see *Materials and Methods* for the positivity criteria). **H**-**I**: OMP-(**H**) specific CD8^+^ cells detected using EO2316-, EO2317 and EO2318 specific tetramers and TAAp-(**I**) specific CD8^+^ cells detected using IL-13RA2-, BIRC5-, and FOXM-tetramers on patients’ PBMC samples *ex-vivo*. For Patient #3, no TAAp-specific data were available. Each dot represents the percentage of tetramer^+^ CD8^+^ cells in each patient and at the indicated week. Negative values are plotted as 0.001. For Patients #1 and #2, only data from the monitoring of the later timepoints (12-24 weeks) were available. **J.** OMP-specific (top) or TAAp-specific (bottom) CD8^+^ cells from PBMC samples *ex vivo* or after IVS.

The substantial T cell responses detected following IVS led us to investigate these responses directly *ex vivo*. Circulating antigen-specific T cells induced post-immunization were quantified without prior *in vitro* expansion. A significant number of EO2317- and EO2318-specific CD8^+^ T cells were detected *ex vivo* using MHC tetramers in three 3 patients, with Patient #1 showing 4.7% and 2.7% of EO2317- and EO2318-specific CD8^+^ T cells, respectively. EO2316-specific CD8^+^ T cells were detected only in Patients #1 and #3 (**Fig. S4I**, gating strategy; **Fig. S4J,** representative staining; **Fig. 5H**). *Ex vivo* TAAp-specific T cells were investigated only in Patients #1 and #2. BIRC5-specific CD8^+^ T cells were detected *ex vivo* in both Patients, reaching 3.3% in Patient #1. FOXM1-specific CD8^+^ T cells were detected *ex vivo* in Patient #1 only, whereas IL-13RA2-specific CD8^+^ T cells were not detectable *ex vivo* (**Fig. 5I).** The potent proliferative capacity of these OMP-specific T cells was underscored when we plotted the number of specific cells obtained *ex vivo* and the corresponding post-IVS levels for Patient #1 (**Fig. 5J**). The memory phenotype of vaccine-expended CD8^+^ T cells was further evaluated *ex vivo* in Patients #1 and #2. We demonstrated that the vast majority (>90%) of these cells were specific effector memory T cells (T_EM_ and T_EMRA_) (representative staining **Fig. 6A, 6B and Fig. S4K**). The functionality of these cells was also evaluated *ex vivo* using IFN-γ ELISpot for Patient #1 and #2 at weeks 44 and 14. We confirmed that the vaccine-specific T cells were not only activated by the individual OMPs, but also cross-reacted with the TAAps (here tested as a pool) (**Fig. S4L**).

**Figure 6.**
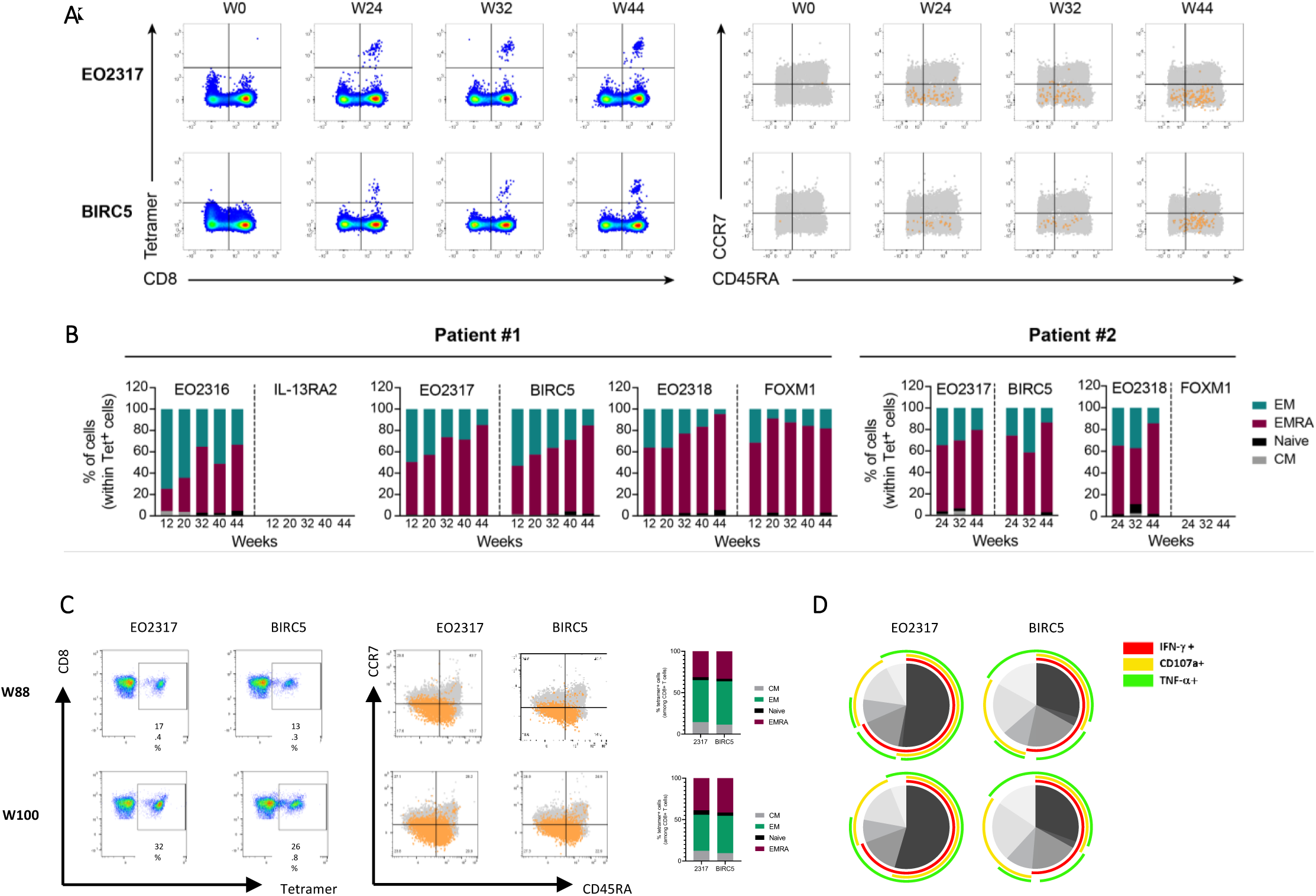
Long-term memory and polyfunctional T cell responses in glioblastoma patients post-vaccination. **A-B.** Vaccine-specific CD8^+^ T cells are effector memory cells Representative dot plots (**A**) for Patient #2 of EO2317- and BIRC5-specific CD8^+^ T cells over time (weeks (W) 0, W24, W32, and W44) based on tetramer staining (left). The memory phenotype (right) was evaluated based on the expression of CCR7 and CD45RA markers. CCR7 and CD45RA expression profiles of tetramer^+^ cells (orange dots) overlaid on CD14/CD19 − cells (grey dots). **B.** Differentiation of subsets of tetramers ^+^ CD8^+^ T cells. Quantification of the memory phenotype of tetramer^+^ CD8^+^ T cells (central memory (CM), naïve, effector memory (EM), and terminally differentiated effector memory (EMRA)) for Patients #1 and #2 are shown as percentages (%). Only time points with more than 20 tetramer+ CD8+ T cells were used for analysis. **C-D.** <u>Durable and long-term response of Patient #1 to vaccination</u>. T cells from Patient #1 were analyzed at weeks 88 and 100 after administration of the first vaccine dose. Flow cytometry analysis (**C**) showing EO2317- and BIRC5-specific T cells (left) and CCR7/CD45RA expression profile (middle) for EO2317- and BIRC5-specific T cells (orange) overlaid with the overall CD8^+^ T cell population (gray population). Central memory, effector memory, naïve, and terminally differentiated effector memory CD8^+^ subsets were quantified for each of the indicated tetramer^+^ populations (right). T cell polyfunctionality is shown in the pie chart (**D**). T cells were expanded in culture with the OMP pool and IL-2 for 12 days and restimulated with either EO2317 or BIRC5 peptide for 6h before being intracellularly stained for IFN-γ, CD107a, and TNF-α. The pie arcs depict the proportion of cells that produce a specific cytokine, and pie slices the proportion of cells co-producing 1 to 3 different cytokines. Pie arc overlap represents polyfunctional cells.

Our clinical protocol includes monthly recall injections, which could potentially lead to T-cell exhaustion. To demonstrate that our approach can sustain a robust T-cell response over an extended period, we analyzed cross-reactive EO2317- and BIRC5-specific T cells in Patient #1 at weeks 88 and 100 post-treatment initiation. Initially, at week 44, EO2317-specific T cells accounted for 3.27% and BIRC5-specific T cells for 1.99% of total circulating CD8^+^ T cells (**Fig. 5H and 5I**). By week 88, these percentages had increased to 17.4% and 13.3%, respectively, and further increased to 32% and 26.8% by week 100 (**Fig. 6C**). This progression demonstrates the substantial and sustained expansion of vaccine-induced lymphocytes over time. Additionally, those long-term induced EO2317-specific CD8^+^ T cells not only maintained their memory phenotype but also their polyfunctionality, resulting in a durable and potent BIRC5-directed response (**Fig. 6C and 6D**).

## Discussion

In the present study, we demonstrated that OMPs, sharing certain degree of homology with TAAps, can be used to generate strong and durable cross-reactive immune responses against cancer cells.

The efficacy of peptide-based immunotherapies relies, in large part, on the properties of the peptides used to activate the immune response against tumor cells. Ideally, these peptides should be highly immunogenic, tumor-specific, and activate a T-cell population that is prevalent in all cancer patients. The quest to identify such peptides remains a formidable challenge. Recent studies indicate that among patients that underwent ICI administration or neoantigen vaccination, long term survivors harbor tumor-specific T-cell clones with predicted cross-reactivity to pathogenic bacteria or viral antigens, suggesting that molecular mimicry, T-cell cross-reactivity, and pre-existing immunity could be key factors for the success of such immunotherapies and by extension could be used to improve peptide-based immunotherapies (40–43). Extending these findings, it has also been suggested that the composition of the gut microbiome can impact checkpoint blockade therapy outcomes (44,45). The human microbiome has the capacity to encode billions of potential antigens, a number which is significantly higher than the whole repertoire of known pathogen-derived antigens; thus, the probability of identifying in humans mimics of TAA-derived peptides with the particular properties we have set here (**Fig. 1**), is certainly high.

For many years, cancer immunotherapies have tried to leverage TCR cross-reactivity to improve anti-tumor CTL responses, primarily using analog or heteroclitic peptides designed to provoke stronger immune responses than the native tumor antigen versions. These peptides, intended to improve MHC-anchor residue interaction or modify TCR contacts, have not consistently translated into effective clinical vaccines (46,47). Given that CDPs naturally shape the T cell repertoire, leveraging their inherent immunogenicity offers a more direct and promising strategy. Our strategy selectively targets OMPs that exhibit improved MHC binding due to amino acid mismatches, while preserving the integrity of central TCR contact points to ensure the TCR can recognize both the OMP and the corresponding TAAp similarly. This decision aligns with recent findings indicating that microbiota-derived peptides with altered TCR contacts may be less effective in triggering cross-reactive CD8 T cells and generating impactful cross-reactive neoantigen CD8 T cells in glioblastoma patients (48,49). The T cell repertoire of each individual is shaped by his/her HLA haplotype and the autoantigens encountered during thymic selection. This repertoire is influenced by various environmental factors, including gut microbiota, which might prime naive T cells and generate a pool of memory T cells (50–52). Our OncoMimics approach is designed to engage this pre-existing T cell pool. Essentially, if tumor cells present TAAps that sufficiently resemble to non-self CDPs, such peptides can be used to reactivate a T cell pool that cross-reacts with TAAps and potential lead the tumor control.

OMPs were selected using a bioinformatic pipeline based on the identification of commensal-derived peptides sharing sequence similarities with known TAAps. The selection process involved a BLAST search that tolerated mismatches at anchor positions and then selecting predicted peptides with high HLA-A2 affinity, efficient predicted cleavage and high prevalence of these OMPs within the human microbiome. This approach yielded a list of 4403 OMPs with anticipated high affinity and cross-reactivity potential. Among those OMP candidates homologous to the analyzed TAAps, we selected the ones with the best score for binding, cleavage and prevalence. Interestingly, we demonstrated for all these OMPs a higher HLA-A2-binding affinities and increased peptide-MHC stability compared to their TAAp counterparts, making them potential candidates to drive strong CTL responses. Their immune potential was further assessed on an A2-DR1 humanized animal model. In this setting, while most OMPs induced stronger TAAp-specific T cell responses compared to the TAAps themselves, few exceptions were noted. For instance, the immunogenic responses to CD22 and FOXM1 were lower when using OMPs compared to when using their corresponding TAAps. This might be attributed to inherent model limitations. Specifically, the potential perception of these human TAAps as foreign peptide, due to amino acid variances with the endogenous mouse equivalent (**Supplementary Table S3**), and/or the absence of these human OMPs in mouse gut microbiota, that could skew to a biased T cell repertoire. Such factors potentially limit our model to fully recapitulate human immune responses, highlighting the need for cautious interpretation of those comparative immunogenicity results.

We pursued our analysis to explore *in vivo* the capacity of these OMPs to trigger functional CTL response against TAAps. We observed that OMPs were indeed able to induce cross-reactive TAAp-specific T cells able to recognize TAAp-loaded tumor cells. We also confirmed in a tumor regression model that OMP72-specific T cells responses triggered by OMP72 were able to induce the regression of CD20 expressing tumor cell. Our findings are in line to other strategies utilizing pathogen-derived peptides similar to TAAps to provoke cross-reactive T-cell responses, addressing the low immunogenicity of natural TAAps and capitalizing on pre-existing high-affinity memory T cells from past infections in line with proposed strategies (53,54). Yet, the success of this pathogen mimic strategy is inherently limited by individual pathogen exposure histories, complicating the development of universally effective treatments. In contrast, our investigation into the widespread presence of OMP-specific T cells in the human population revealed that more than 80% of tested healthy donor PBMCs (for 8 out of 10 OMPs) contained such cells. Furthermore, these OMP-specific T cells, when expanded *in vitro*, demonstrated reactivity against their corresponding TAAps, as shown by tetramer staining analysis. More importantly, those expended cells exhibited comparable cytotoxicity against TAP-deficient T2 cells presenting either the same OMP or its corresponding TAAp, even at low E:T ratios. This evidence supports the potential of utilizing OMPs to engage pre-existing T-cell repertoires, offering a promising avenue for developing broadly applicable, off-the-shelf immunotherapeutic strategies.

Building upon these promising findings, specific OMPs have been utilized to develop off-the-shelf immunotherapeutic vaccines, currently undergoing clinical evaluation. EO2401 is a cancer vaccine that includes three OMPs (EO2316/OMP16, EO2317/OMP17, and EO2318/OMP18) that mimic TAAps (IL-13RA2, BIRC5, and FOXM1) and a universal HLA-DR class II peptide derived from telomerase to support CTL cells in combination with the PD-1 inhibitor nivolumab and anti-VEGF bevacizumab. This vaccine is currently being evaluated in a multicenter phase Ib/IIa clinical trial (EOGBM1-18, NCT04116658) in patients with recurrent glioblastoma. The selection of these primary targets (IL-13RA2, BIRC5, and FOXM1) in GBM approach is supported by prior scientific or clinical data validating these tumors antigens as well express TAA in tumor cells of GBM patients (55–57). Initial immunomonitoring data obtained from blood samples of three patients included in the first cohort demonstrated that this approach is unique among other peptide-based therapies in its ability to generate robust CD8+ T cell responses. Specifically, a strong immune response against OMPs and targeted TAAps was induced in all three patients, with *ex vivo* frequencies of tumor-specific systemic T cells reaching up to 1% of the total CD8+ T cell subset. In these patients, we observed distinct patterns of immune responses among the OMP/TAAp pairs: EO2317/BIRC5 and EO2318/FOXM1 elicited higher levels of cross-reactive T cell responses, whereas EO2316/IL-13RA2 demonstrated a lower capacity to trigger cross-reactive T cells. Therefore, the effectiveness of each OMP/TAAp pair is determined by factors beyond the mere peptide-MHC binding affinity. Notably, the intrinsic TCR cross-reactivity rules for each peptide pair, along with variations in individual gut microbiota composition, could significantly influence their effectiveness.

For Patient #1, administration of OMPs led to a significant expansion of circulating TAAp-specific CD8+ T cells (>20% for BIRC5-specific T cells). This remarkable expansion surpasses the typical outcomes observed with classical vaccination approaches, for which the frequency of circulating tumor antigen-specific CD8+ T cells is rarely detectable *ex vivo* by MHC-tetramer (58). These findings demonstrate robust and sustained responses, with *ex vivo* T cell percentages in the single- or even double-digit range. Importantly for therapeutic considerations, OMPs elicited persistent tumor antigen-specific T cell responses in Patient #1, extending for almost 2 years of treatment, while maintaining their cross-reactivity and polyfunctional properties. Notably, these levels exceed in the long term those reported for CAR-T or TCR-T therapies, for which the numerous counts of cells initially injected rapidly declines over time (59,60). The rapid and sustained immune responses observed in the three patients as early as four weeks after the first administration suggest that our approach targets a highly proliferative T cell population, which we hypothesized to be a pre-existing memory cell population. This observation is reinforced by the level of immune response and the large number of circulating memory T cells (based on CD45RA/CCR7 markers) observed soon after vaccination, which demonstrates the robust *in vivo* expansion of effector memory CD8^+^ T cells. These observations strongly support the proposed strategy that memory T cells activated upon OMP vaccination are initially generated through the exposure of gut T cells to commensal peptides. However, whether these OMP-specific T cells originate from a pre-primed, gut-derived memory T cells pool remains to be determined. In line with these results, the IVS experiments demonstrated that only OMPs efficiently induced the proliferation of TAAp-specific T cells in PBMCs isolated from Patient #1. Functional analysis of these post-IVS T cells showed that they were not only highly proliferative, but also polyfunctional, with most of these TAAp-specific T cells displaying an effector memory phenotype (T_EM_ and T_EMRA_). Our study primarily investigated the efficacy of OMPs in eliciting OMP-/TAA-specific cross-reactive T cells in the peripheral blood. Consistent with studies showing that antigen-specific T cells can migrate from peripheral blood and infiltrate brain tumors after peptide-based immunotherapy (49,61,62), preliminary data from patient-derived relapsed tissues show CD8+ T cell recruitment within tumors following EO2401 treatment (63). To conclusively establish the specificity and migration patterns of vaccine-induced T cells, further in-depth analyses involving TIL isolation and subsequent tetramer staining or single-cell TCR profiling are needed. Corroborating our approach, we note that a recent study demonstrated that glioblastoma CD4^+^ TILs can recognize a diverse array of peptides, including peptides derived from commensal gut microbiota (64).

In summary, we have shown that commensal bacteria-derived peptides, with homology to naturally occurring TAAps, when carefully chosen through a combined bioinformatic and lab-assay pipeline, can be used to generate rapid and long-lasting immune CD8^+^ T cell responses at high frequencies. We applied this approach for the first time to vaccinate patients with glioblastoma, a low mutation burden tumor for which ICIs trials have not shown any clinical benefit. Extremely potent and long-lasting immune responses were observed in the first three patients. Although follow-up is required to confirm these conclusions, we observed that two of the three patients survived beyond 1 year. This outcome, although preliminary, offers a cautiously optimistic perspective in the context of this challenging clinical condition (65).

## Acknowledgments

We would like to thank the atients and their families who participated in Cohort 1 of the EOGBM1-18 study (NCT04116658). We would like to express our sincere gratitude to Professor Pedro Romero of the University of Lausanne, Professor David Klatzmann, Professor of Immunology at Sorbonne Université, and Head of Biotherapy at Pitié-Salpêtrière Hospital for their essential comments, suggestions, and manuscript review. Their expertise advanced significantly and shaped the project. We thank Muriel Mas, Jean-Michel Paillarse, and Jan Fagerberg, whose pivotal roles were instrumental in steering the clinical development of this project. We thank all Enterome employees for their dedication that enabled this study. We also thank Rachel Morra and Guillame Bayre for their assistance with editing and proofreading. Figures were created using Biorender.com.

## Author contributions

JGM, CB^1^ and LC conceived and designed the study, drafted the original manuscript, and supervised the project. AT, AM, JMC, GK, and JGM performed experiments. AT, JMC, GK, AM, LA^1^, LA^2^, JK, TM, AG, CPO, CC, MB, DB, CV, JM, JN, KL, LB, MM, OA, OJ, FS, LC, and JGM conducted formal analyses. CB^2^, CG^1^, FS, and JGM were responsible for the data curation. AT, JMC, GK, AM, LA^1^, JK, FS, and JGM contributed to the methodology development. OA, OJ, CG^2^, LC, and JGM administered the project. AM, CG^2,^ AI, MV, FG, AS, GT, MCB, IM, UH, WW, and DAR contributed the resources. AT, JMC, GK, AM, LA^1^, JK, CG^2^ and JGM verified the replication and reproducibility of the results and created the visualization and data presentation. AT, JMC, AM, LA^1^, JK, GC, CG^1^, FS, CG^2^, LC and JGM reviewed and edited the manuscript. Christophe Bonny (CB^1^); Coline Billerey (CB^2^); Camille Gaal (CG^1^) and Cécile Gouttefangeas (CG^2^) Lucie Aubergeon (LA^1^), and Ludivine Amable (LA^2^). All authors read and approved the final manuscript.

## Supplementary figures

**Supplementary Fig. S1:**
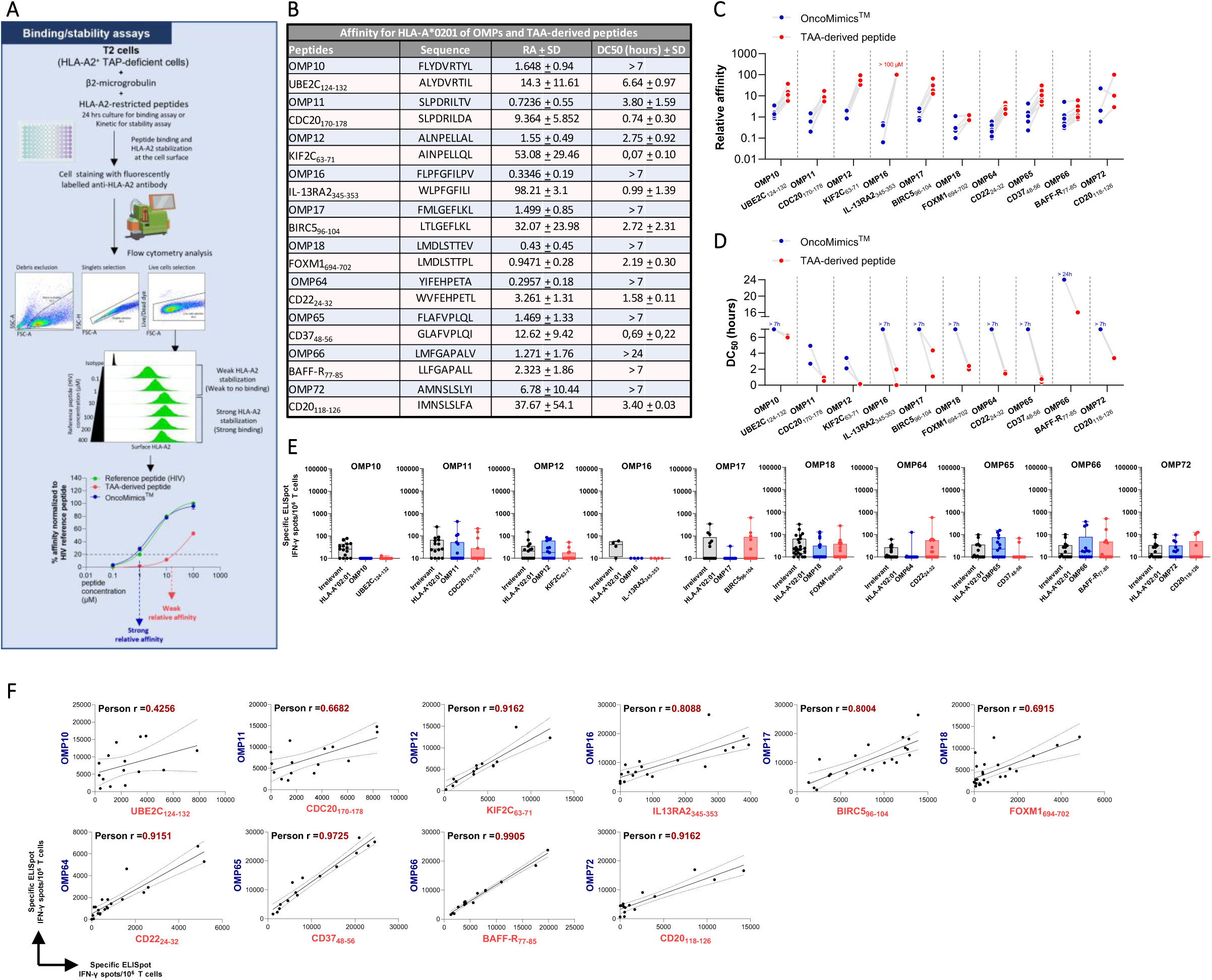
**A.** <u>Schematic representation of HLA-A2 T2 binding and stability assay.</u> **B.** Relative affinity and dissociation complex values calculated for each peptide. The relative affinity (RA) is calculated as the concentration of each peptide normalized to the concentration of the reference peptide, that induces 20% of stabilized HLA-A2. The dissociation complex (DC50) represents the half-life of the peptide/HLA-A2 complex. SD=standard deviation. **<u>C, D.</u>** <u>Relative affinity and dissociation complex values for each independent experiment</u>. **<u>E.</u>** <u>OMP- and TAAp-induced TAA-specific IFN-γ responses.</u> Comparison of OMP efficacy in inducing a TAA-specific response to that of their TAA counterparts in A2/DR1 humanized mice, assessed on splenocytes from naïve control mice with the indicated peptides by IFN-γ ELISpot assay. **<u>F.</u>** <u>Correlation analysis between IFN-</u>γ <u>ELISpot response to OMPs and their TAAp counterparts in mice immunized with OMPs.</u> The analysis was performed using a Pearson correlation coefficient assuming a Gaussian distribution.

**Supplementary Fig. S2:**
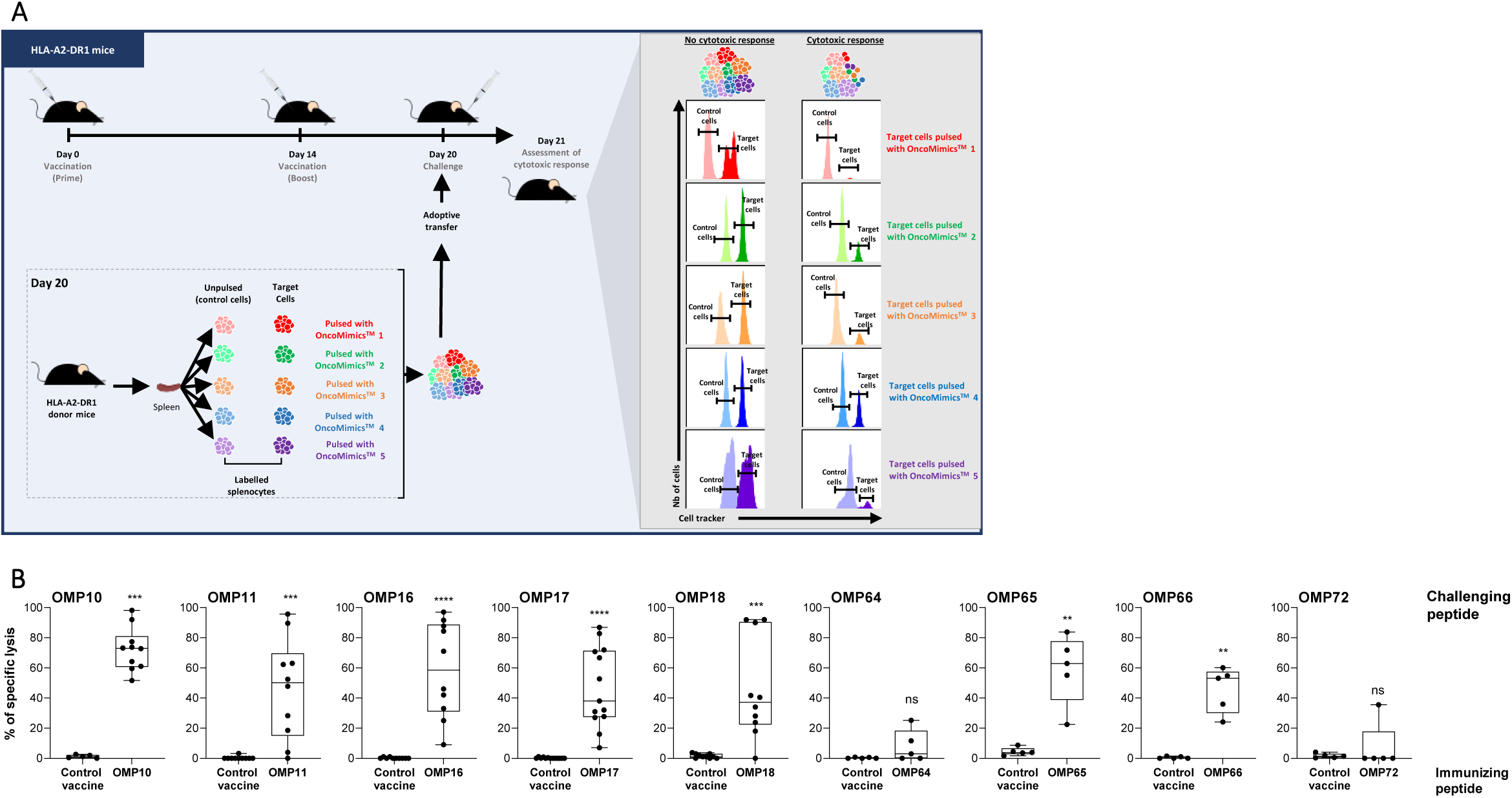
**A.** <u>Schematic representation of the experimental setup employed to assess the *in vivo* T cell cytotoxicity elicited by each OMP.</u> HLA-A2-DR1 mice were vaccinated with OMPs and challenged post-vaccination with syngeneic splenocytes labeled with multiple cell tracking dyes (Cell Trace Blue, Cell Trace Violet, CFSE, Cell Trace Yellow and Cell Trace Far Red). Target cells were labeled with high concentrations of dyes (intense colors) and pulsed with individual OMPs before being mixed altogether and with unpulsed dim control cells (light colors on schematic) at equal ratio. The cell suspension was adoptively transferred and the cytotoxic responses against up to 5 OMPs could be analyzed 20 h post-challenge. **B.** <u>In vivo cytolytic activity elicited by individual OMPs.</u> Percentages of *in vivo* specific lysis of splenocytes pulsed with the indicated pool of OMPs after immunization with the indicated peptide pool.

**Supplementary Fig. S3:**
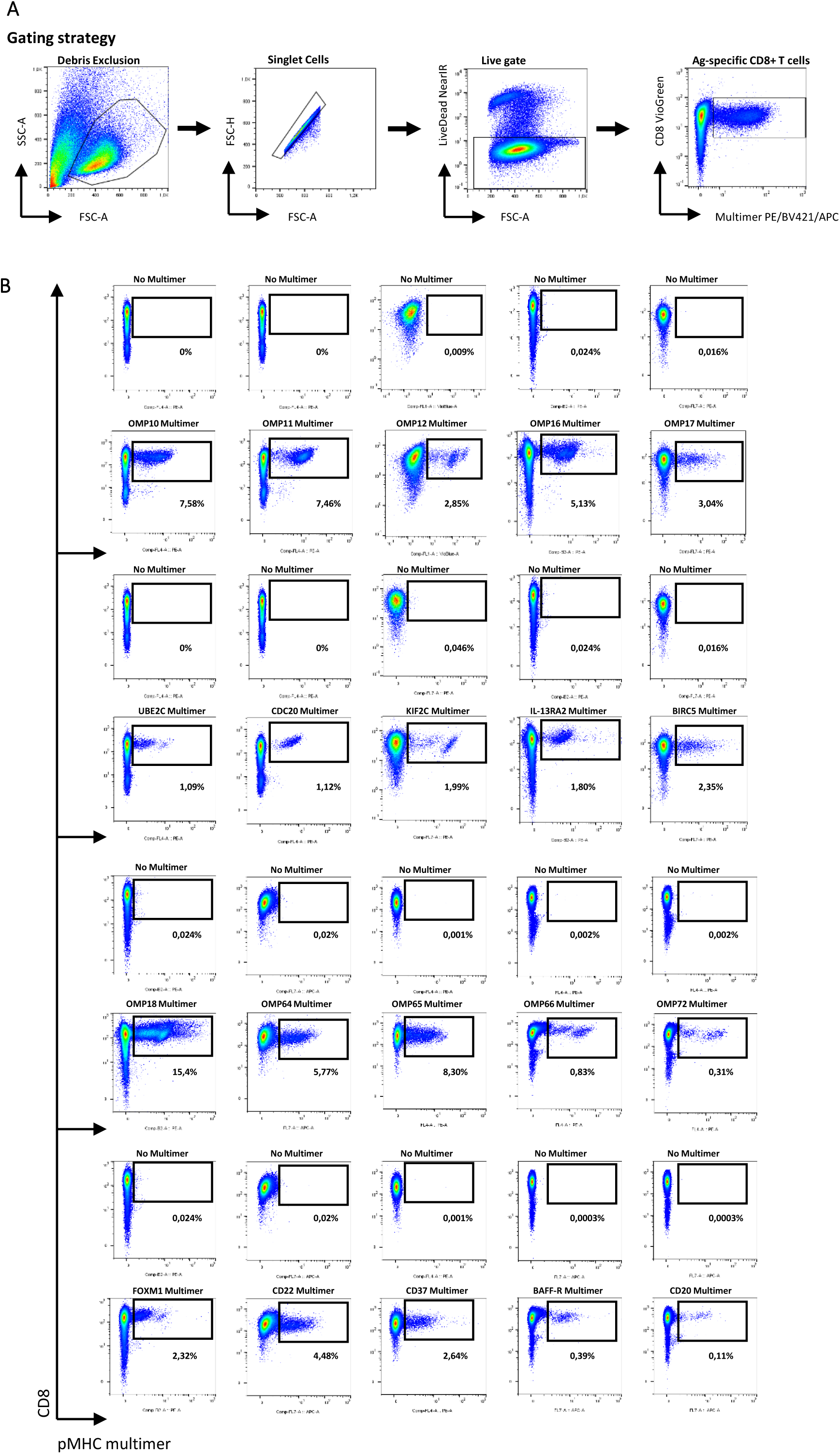
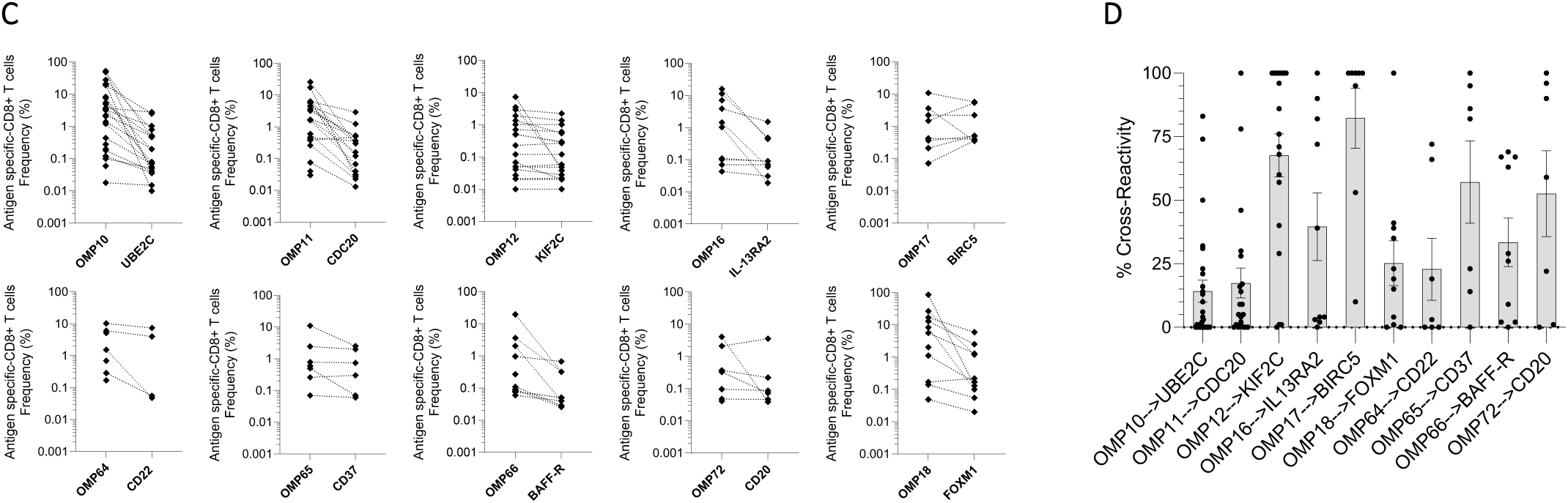
**A.** <u>Gating strategy and representative results of a pMHC tetramer staining by flow cytometry.</u> First, the lymphocyte population was delineated based on its forward scatter area (FSC-A) versus side scatter area (SSC-A) properties to exclude cell debris. Then, a gate on diagonally clustered singlets was created by plotting FSC-A versus FSC-height (FSC-H). Live cells were gated in LIVE/DEAD marker negative versus FSC-A. Next, the gate was set on the CD8+ cells that are positive in the tetramer channel. This gate was used to identify the CD8+ tetramer+ cells (antigen-specific CD8+ T cells). **B.** <u>Detection of antigen-specific CD8+ T cell for OMPs and TAAp-derived peptides using conjugated pMHC mulimers after expansion of PBMCs with OMPs.</u> Representative staining data are shown for each peptide. pMHC multimer stainings performed with the indicated conjugated pMHC multimer for each OMP and TAAp. Conditions without pMHC multimer were used as negative control to define the positive pMHC multimer gate and named no multimer. Percentage of tetramer+ in CD8+ T cells is indicated in the dotplots. **C.** <u>Induction of OMP-specific cross-reactive CD8+ T cells against TAAps.</u> Frequencies of antigen-specific CD8+ T cell clones for OMP and TAAp were evaluated in healthy donors after three rounds of IVS with OMPs. Each dot represents the frequency obtained for each individual donor (n=7 to 27). **D**. <u>Percentage of cross-reactivity between OMP and TAAp in HDs.</u> Each dot represents cross-reactivity frequency determined as specified in Material and Methods for each individual donor after three rounds of IVS with OMPs (n=7 to 27). Data are represented as mean ± SEM.

**Supplementary Fig. S4:**
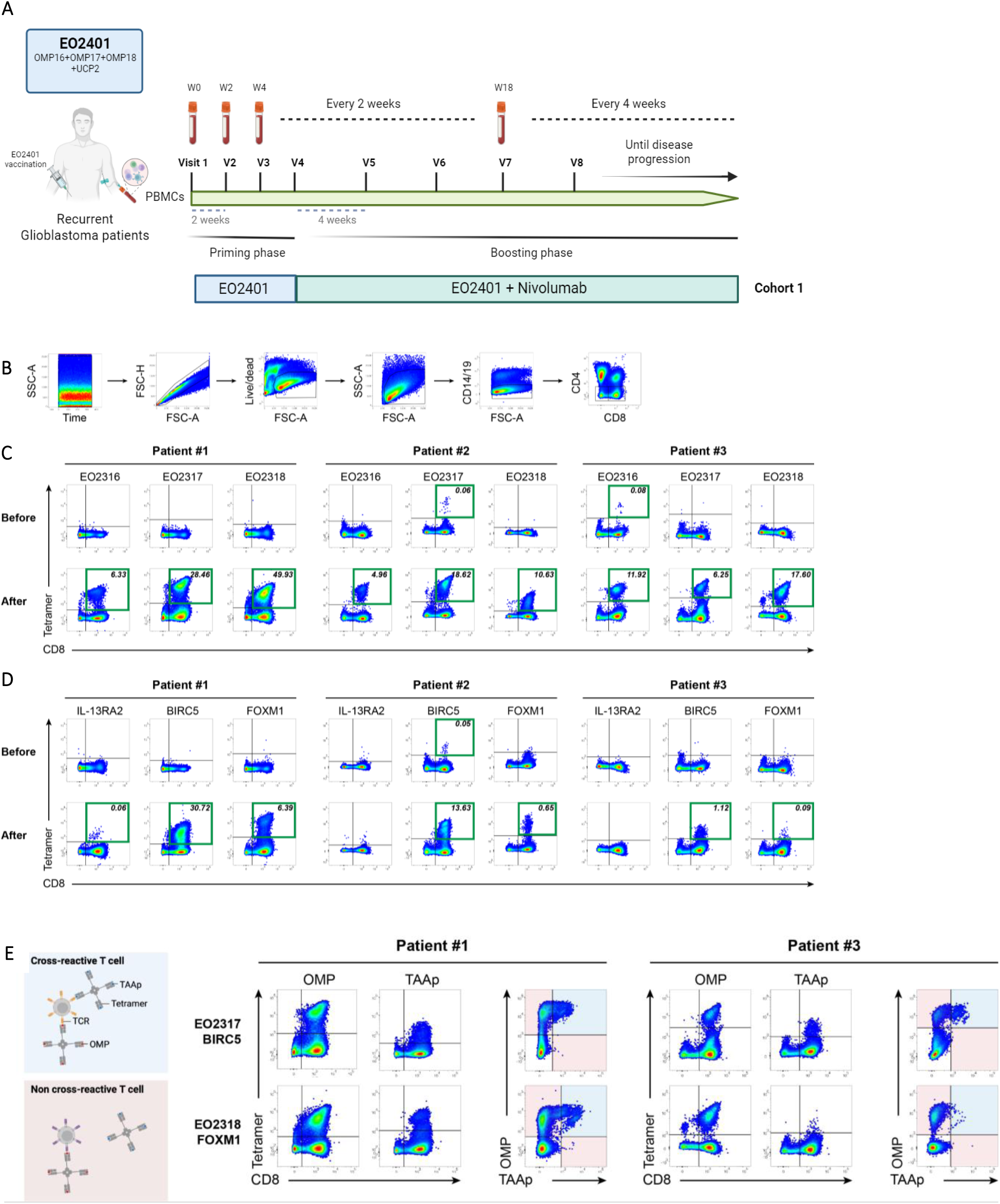
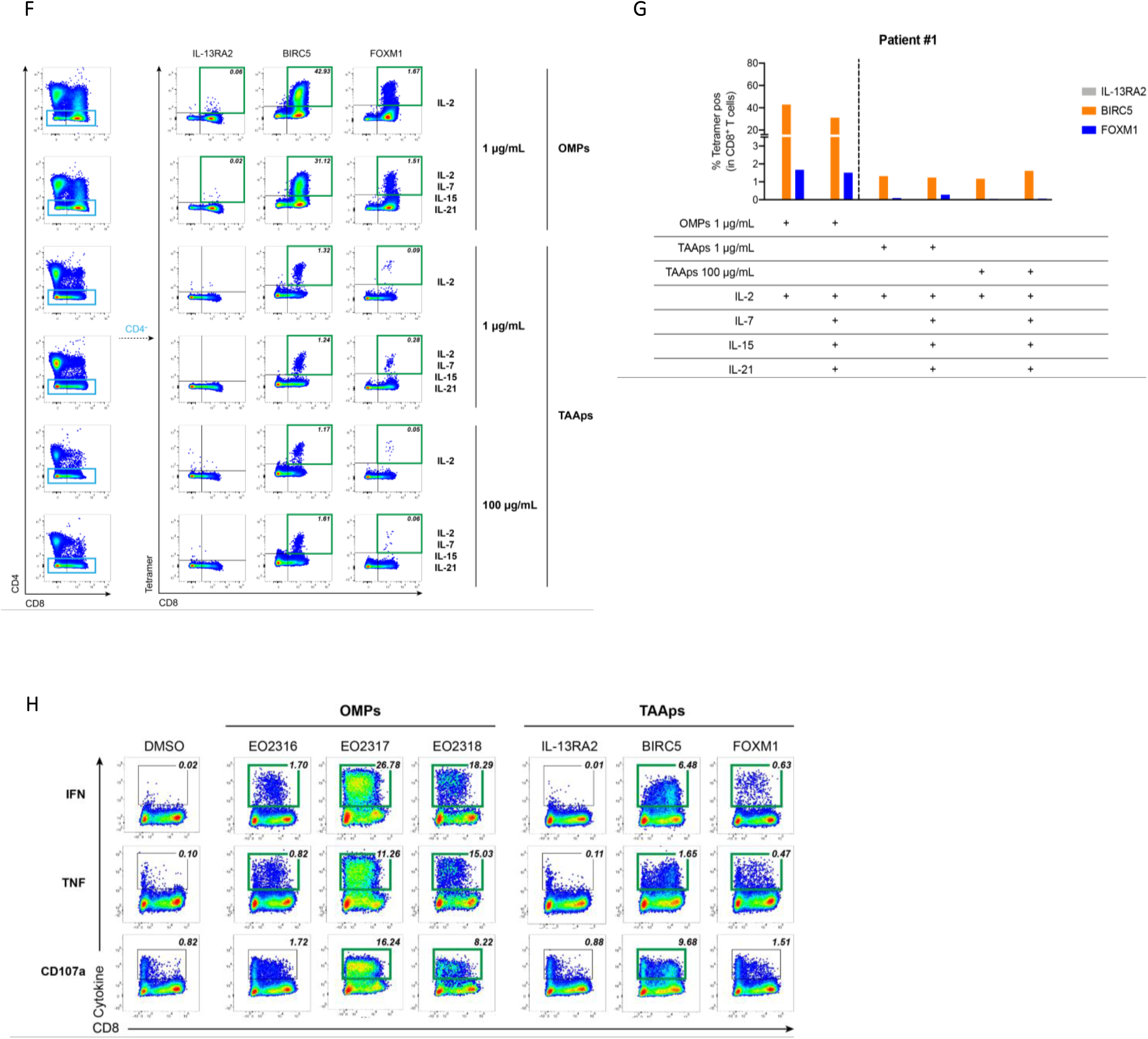
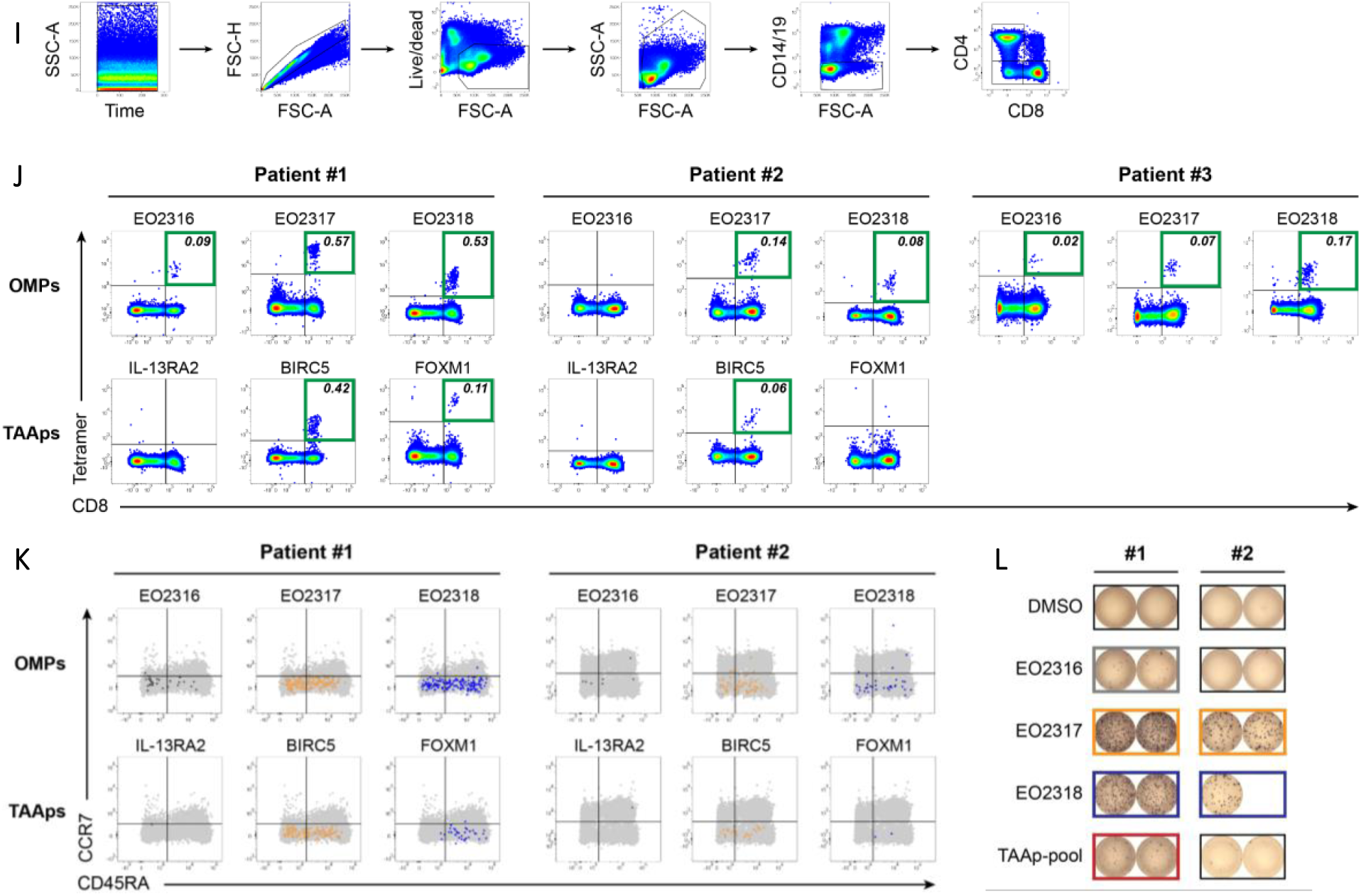
**A.** <u>Schematic overview of EOGBM1-18 clinical trial (Cohort 1)</u>. Treatment schedules and sampling of EOGBM1-18 clinical trial. Patients with recurrent glioblastoma receive 4 doses of EO2401 every two weeks followed by EO2401 (every 4 weeks) + nivolumab, as shown in schematics. PBMCs are isolated at multiple time points (either every two or four weeks) and used for immune response monitoring. **B.** <u>Gating strategy used in tetramer staining after IVS</u>. Briefly, doublets, dead cells, CD14^+^ and CD19^+^ cells were excluded. Cells were then gated on CD4^-,^ and tetramer^+^ cells were counted in CD8^+^ cells. **C**-**D.** <u>Representative dotplots of tetramer staining</u> for Patient #1 (left panel), #2 (middle panel) and #3 (right panel). Staining with OMPs (EO2316, EO2317 and EO2318) and TAAps (IL-13RA2, BIRC5, FOXM1) are shown in **C** and **D**, respectively. Percentage (%) of antigen-specific CD8^+^ cells is shown for samples which were considered positive (highlighted with a green square) (see *Materials and Methods* for positivity criteria). Upper and lower row show before and after vaccination. **E.** <u>Cross-reactivity between OMP and TAAp</u> is shown for EO2317 (BIRC5) and EO2318 (FOXM1). For this, OMPs and TAAps were included in the same staining mix and double-positive cells were investigated. Representative dot plots for Patients #1 and #3 are shown. Non- and cross-reactive CD8+ T cells are highlighted in red and blue, respectively. **F-G**. Expansion of TAAp-specific CD8^+^ T cells after *in vitro* stimulation with OMPs- or TAAps-<u>pool</u>. PBMCs from Patient #1 were either stimulated with 1 µg/mL of OMPs or TAAps or 100 µg/mL of TAAps in the presence of either IL-2 alone or IL-2, IL-7, IL-15 and IL-21. After 12 days, cells were collected and TAAp tetramer staining was performed. <u>Representative dotplots of tetramer staining</u> for Patient #1 and their quantification are shown in **F** and **G**, respectively. Percentage (%) of antigen-specific CD8^+^ cells is shown for samples which were considered positive (highlighted with a green square) (see *Materials and Methods* for positivity criteria). **H.** <u>Representative dot plots of ICS</u> for IFN-γ, TNF and CD107a for cells stimulated with the respective peptide shown on top. Gating strategy shown in A was applied and CD4^-^ T cells are shown in the plots. Percentage (%) of activation marker positive cells within CD4^-^ T cells is shown in dot plots and positive reactions (see *Materials and Methods* for criteria) are highlighted with green squares. **I.** <u>Gating strategy used in tetramer staining *ex vivo*.</u> Same strategy as described for **B** was applied. **J.** <u>Representative dot plots</u> of tetramer staining for Patients #1 (left panel), #2 (middle panel) and #3 (right panel) after vaccination. Staining with OMPs (EO2316, EO2317 and EO2318) and TAAps (IL-13RA2, BIRC5, FOXM1) are shown in upper and lower row, respectively. Percentage (%) of antigen-specific CD8^+^ cells is shown for samples which were considered positive (highlighted with a green square) (see *Materials and Methods* for criteria). **K.** <u>Memory phenotype analysis</u> of tetramer^+^ cells detected in Patients #1 and #2. Patient #3 was not analyzed. Memory phenotype was evaluated based on CCR7 and CD45RA marker expression. Tetramer^+^ cells (colored dots) are overlayed in CCR7 against CD45RA in CD14^-^/CD19^-^ cells (grey dots). The differentiation subsets central memory (CM, CCR7+/CD45RA-), effector-memory (EM, CCR7-/CD45RA-), naïve (CCR7+/CD45RA+) and terminally differentiated effector memory (EMRA, CCR7-/CD45RA+) are depicted. **L.** <u>Ex-vivo IFN-γ ELISpot</u> for Patients #1 (left panel) and #2 (right panel). Positive wells are highlighted in color.

## Supplementary Tables

**Supplementary Table 1:** <u>List of the 224 TAA-derived peptides used for the analysis</u>. The tables provide the following information for each antigen. The UniProt accession number (<u>TAA_uniprot_ID</u>) and the gene name of the antigen (<u>TAA_uniprot_GN</u>), the antigen-derived peptide sequence (<u>TAAp_sequence)</u>; its position in the protein sequence (<u>TAAp_position)</u>, the length of the peptide (<u>TAAp_length)</u>, the antigen name encoding human gene and/or the parent protein (<u>TAA_name)</u>; the common full gene name (<u>TAA_common_name);</u> the type of tumor-associated antigen, either Shared Antigen, Unique Antigen, B cell marker or Unclassified) <u>(TAA_tumor);</u> the classification, either Alternative ORF, Differentiation, Overexpressed, Shared tumor Antigen, Substitution Mutation, Unclassified or N/A) <u>(TAA_classification);</u> the PMID reference article <u>(PMID_reference)</u>

**Supplementary Table 2:** <u>Data table containing the results of the commensal-derived peptide research from the 224 TAAp.</u> The table output consists of the following columns. The UniProt accession number (<u>TAA_uniprot_ID</u>) and the gene name of the antigen (<u>TAA_uniprot_GN</u>); the antigen name encoding human gene and/or the parent protein (<u>TAA_name)</u>; the antigen-derived peptide sequence in amino acids (<u>TAAp_sequence)</u>; the length of the peptide (<u>peptide_length); its rank and predictive affinity in nM with NetMHCpan 3.0</u> (1) (rank_TAAp and predicted_affinity_TAAp_nM); the commensal-derived peptide sequence in amino acids (commensal_derived_peptide_sequence); <u>its rank and predictive affinity in nM from NetMHCpan 3.0</u> (rank_commensal_derived_peptideand predicted_affinity_commensal_derived_peptide_nM<u>) with the associated MHC class I allele (</u>allele_name) ; the blast match between the TAAp and the commensal-derived peptide (match); the associated percentage of coverage and identity (coverage_percentage and identity_percentage); the number and the positions of mismatches between both sequences, the amino acids equivalences, based on the PAM30 matrix are considered as mismatches (mismatches_number_no_equivalence and mismatches_positions_no_equivalence); the comparison of the predicted affinities, “Yes” if the commensal-derived peptide has a stronger predicted affinity than the associated TAAp, “No” if not (better_predicted_affinity_of_commensal_derived_peptide_compare_to_TAAp); the cleavage prediction score of the commensal-derived peptide (cleavage_prediction_score:Netchop3.0)(2); the commensal protein name (commensal_protein_name) and amino acid sequence (commensal_protein_sequence); the commensal bacteria annotations (phylum, genus, species and KO); the predicted commensal protein localization, either cytoplasmic, transmembrane or secreted (commensal_protein_localisation); the commensal protein prevalence, defined as the percentage of samples from the human gut microbiome catalogue cohort where the protein is present (commensal_protein_prevalence); the commensal-derived peptide prevalence, defined as the percentage of samples from the human gut microbiome catalogue cohort where at least one of the commensal proteins containing the commensal-derived peptide is present. The commensal-derived peptide prevalence is equal to or greater than the commensal protein prevalence (commensal_derived_peptide_prevalence), which is defined as the number of unique commensal proteins containing the associated commensal-derived peptide (commensal_protein_diversity).

**Supplementary Table 3:**
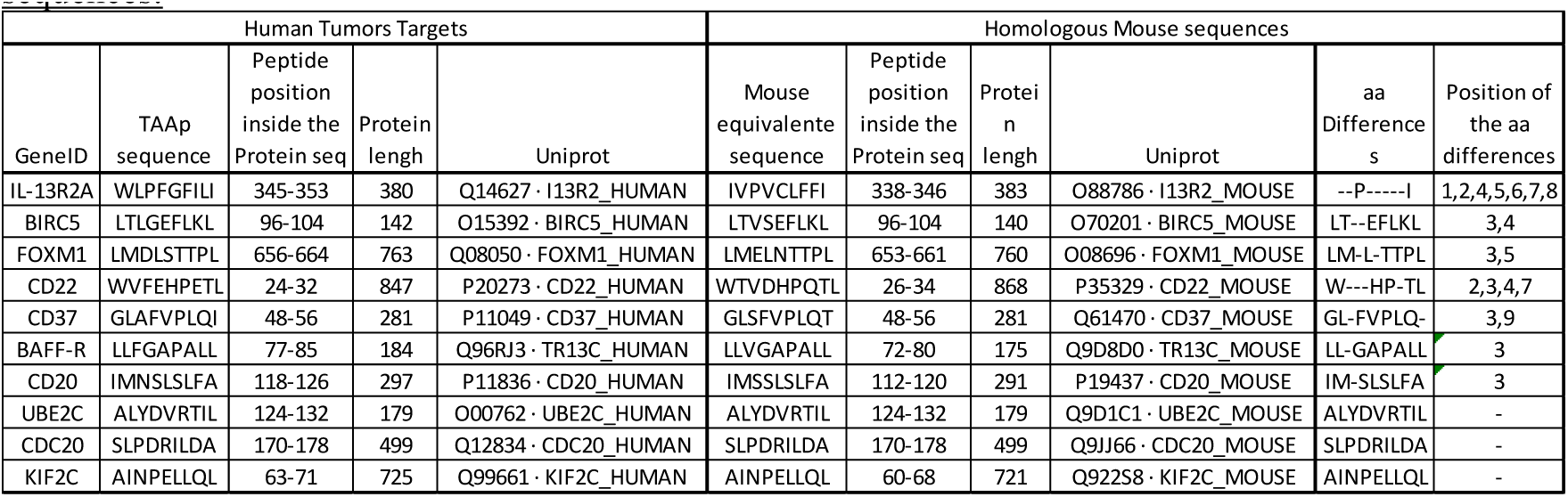
Characteristics of the ten selected TAAps in this study and their homologous mouse sequences: For the human tumor targets, we display the Gene ID of the targeted TAA, the amino acid (aa) sequences of the TAAps, the peptide positions of the TAAps inside the TAA protein sequences, the length in aa of the TAA proteins, and the UniProt accession numbers of these TAAs. For the homologous mouse sequences, we displayed the aa sequences of the mouse equivalent sequence for the given TAAp, the peptide positions of these sequences inside the mouse equivalent protein sequences, the length in aa of the mouse proteins, and the UniProt accession numbers of the mouse proteins. Amino acid substitutions are indicated by a “-,” and similar positions are also shown. The number of mismatch positions between the human and mouse peptides is displayed in the column titled ‘Position of the Amino Acid Differences.

## Supplementary Methods

### In silico OncoMimics^™^ selection

The identification process for the OMP is shown in **Fig. 1A**. A comprehensive list of TAAs was compiled from public databases (TANTIGEN 2.0 (3) and CAPED databases (https://caped.icp.ucl.ac.be/) and CAD v1.0 (4)) and enriched with TAAs reported in prior studies. Following the compilation of TAAs, we assessed data from resources including : Expression Atlas (5), GTEx (6), GEPIA (7), The Human Protein Atlas (8), HLA Ligand Atlas (9) and SysteMHC Atlas (10,11) to curate a list of TAAs with cancer-specific expression. This process yielded 113 unique TAAs. Subsequently, we selected 224 HLA-A2-specific tumor peptides derived from these TAAs and characterized as HLA-A2 binders, naturally presented peptides or as T cell epitopes shown to elicit *in vivo* tumor rejection. Detailed references and peptide information are provided in **Supplementary Table S1**. For the identification of CDPs mimicking these TAAps, a sequence similarity search was conducted against a catalogue of human gut microbiome proteins based on a collection of several million bacterial genes (12). This search pinpointed CDPs candidates with differences allowed only at anchor positions (1, 2, 8, and 9 or 1, 2, 9, and 10, for the 9- and 10-mer peptides respectively). These CDPs were then assessed for their HLA binding affinity, proteasomal processing scores, and prevalence within the gut microbiome (calculated from the associated commensal protein abundance in the human gut microbiome catalogue), leading to a ranking score based on these criteria and the minimal number of mismatches, detailed in **Supplementary Table S2**. The ten OMPs described in this article were selected based on their high scores across all previously mentioned criteria and their mimicry of key tumor antigens (IL-13RA2, BIRC5, FOXM1, UBE2C, CDC20, KIF2C, CD22, CD37, BAFF-R, and CD20).

### *In vivo* cytotoxicity

To test the cytotoxic function of CD8^+^ T cells that were activated following vaccination, immunized A2/DR1 mice were challenged 6 days post-boost immunization with syngeneic splenocytes pulsed with either a mix of peptides or individual peptides. To do so, unimmunized A2/DR1 donor mice were euthanized and a suspension of syngeneic splenocytes was prepared. To determine the individual contribution of each peptide from an immunization mix, the donor cell suspension was split into as many populations as needed to have control and target cells for each peptide for analysis and labeling with cell tracking dye with different fluorescence properties (Cell Trace Blue, Cell Trace Violet, CFSE, Cell Trace Yellow, Cell Trace Far Red, BioLegend, and ThermoFisher Scientific, **Fig. S3A**), allowing the individual tracking of each population. The bright populations used as target cells were pulsed for 2 h at 37°C with a peptide pool or a specific peptide at a final concentration of 100µM before being mixed at an equal ratio with the other populations. Cells were injected intravenously into immunized and naïve mice and the *in vivo* cytotoxic response was assessed 20 h post-injection. Antigen-specific lysis was calculated as follows:

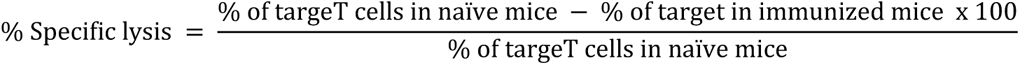

### pMHC tetramer gating strategy

PBMCs were sequentially gated following the hierarchy time parameter histograms: cells that were nonlinearly acquired over time were removed from the analyses. FSC-A vs. FSC-H dot plot: gate on singlets. FSC-A vs. live/dead dot plot: gate on living cells. CD8 VioGreen vs. tetramer (PE/APC/BV421): gates on CD8^+^ and CD8^+^tetramer^+^ cells.

### Trial design and treatment

An ongoing multicenter phase 1/2 trial (NCT04116658) investigated EO2401 (300 µg/peptide, every 2 weeks × 4, then every 4 weeks) and EO2401 with nivolumab (3 mg/kg every 2 weeks) in consenting patients (HLA-A2, KPS ≥70, dexamethasone ≤ 2 mg/day within 14 days before the study, normal organ function, no contraindications) with glioblastoma at first progression/recurrence after surgery and adjuvant radiotherapy/temozolomide. Treatment is administered until toxicity or tumor progression using the iRANO criteria (13). EO2401 is a therapeutic peptide vaccine composed of three HLA-A*02:01 commensal-derived peptides mimicking cytotoxic CD8^+^ T cell epitopes from the TAAs interleukin-13 receptor alpha-2 (IL-13RA2), baculoviral inhibitor of apoptosis repeat-containing 5 (BIRC5), also called survivin, and forkhead box M1 (FOXM1), combined with the CD4^+^ T cell helper peptide universal cancer peptide 2 (UCP2). The peptide mix, that is, the drug product, was emulsified with the adjuvant Montanide ISA 51 VG to obtain a water-in-oil emulsion before subcutaneous administration. Blood collection, standardized PBMCs isolation, and freezing were performed at baseline and then every two or four weeks. (**Fig. S5A**). This study was approved by the ethical review boards of all participating institutions and all participants provided informed consent before enrollment in the study.

### Key resources table

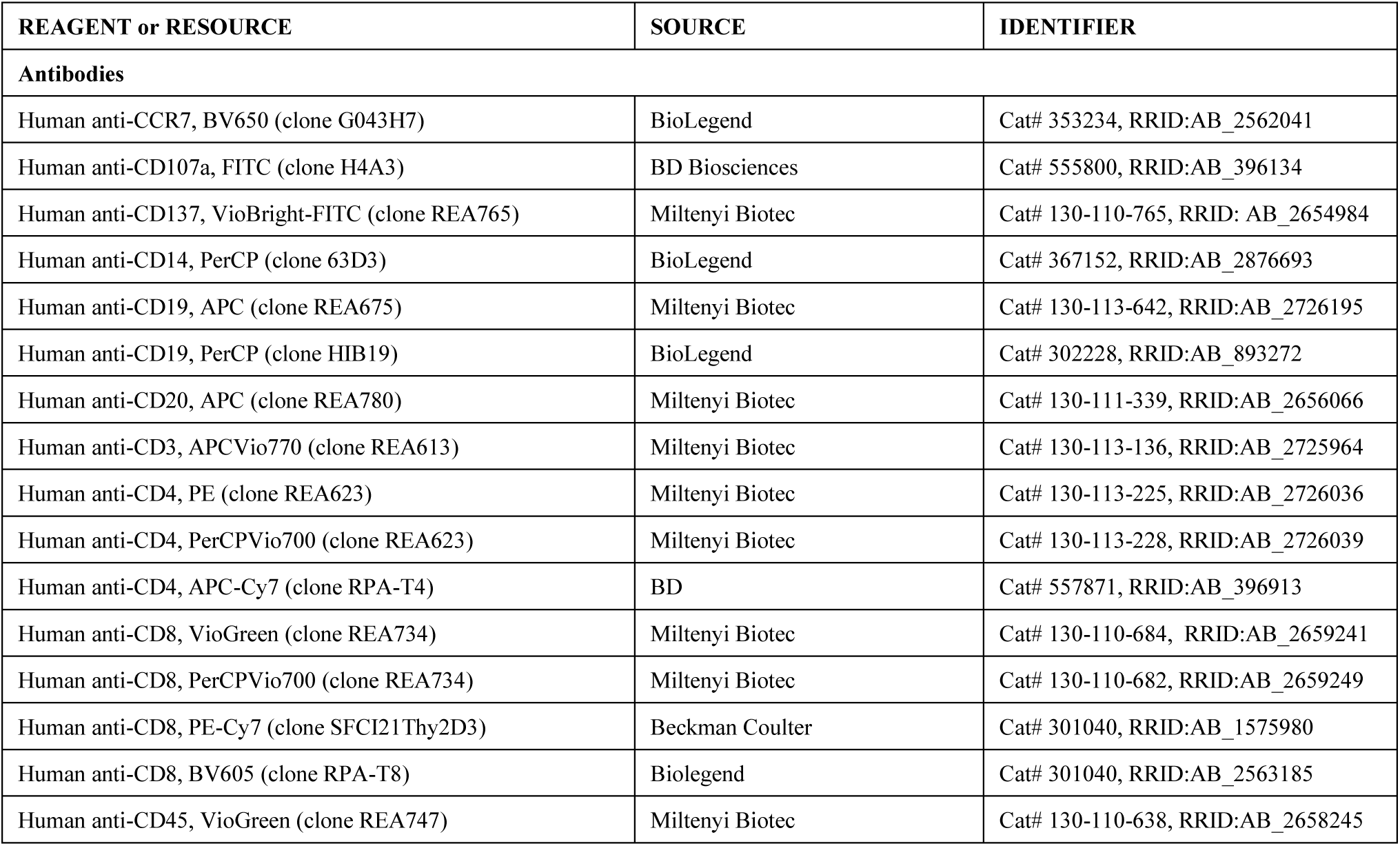
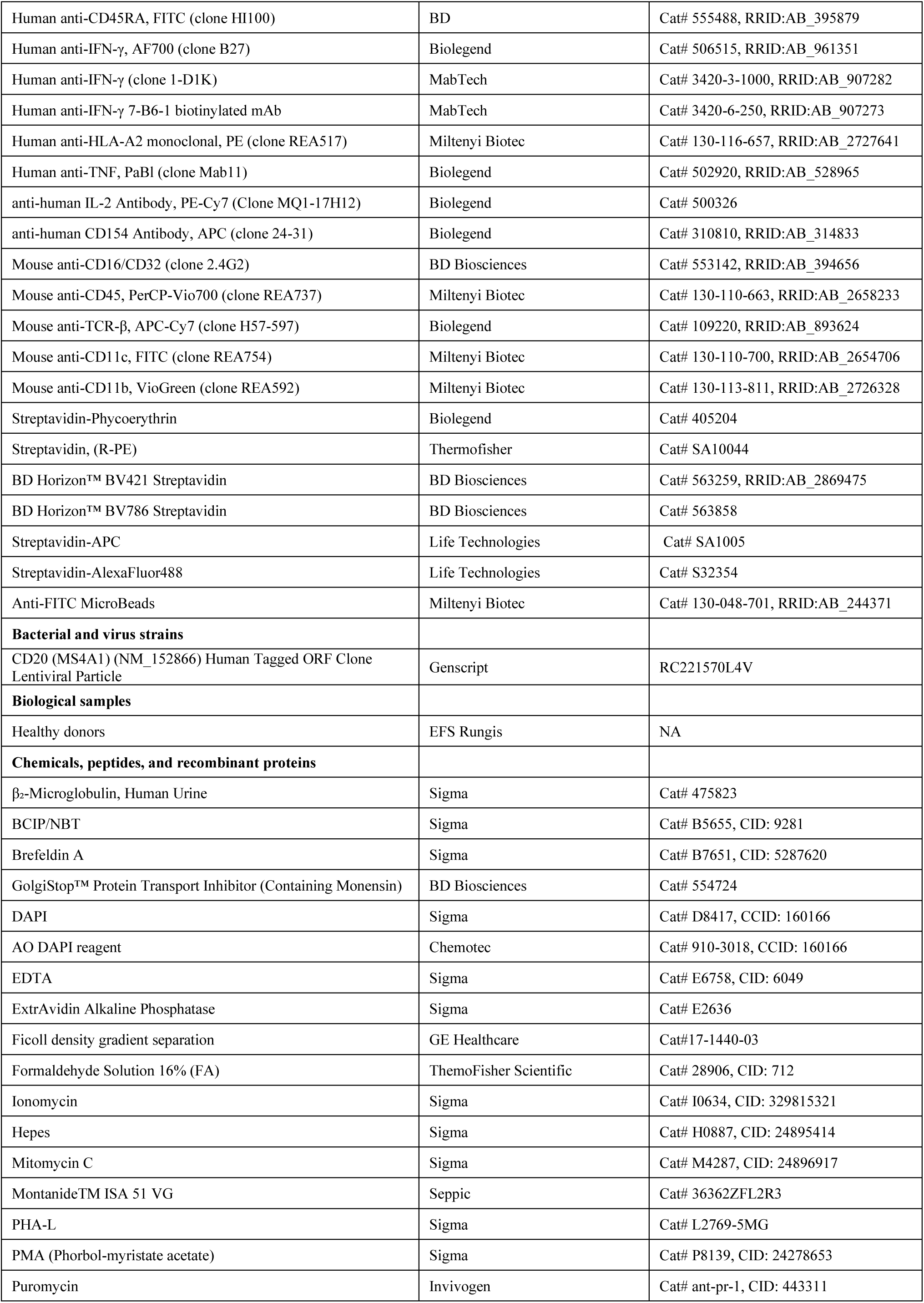
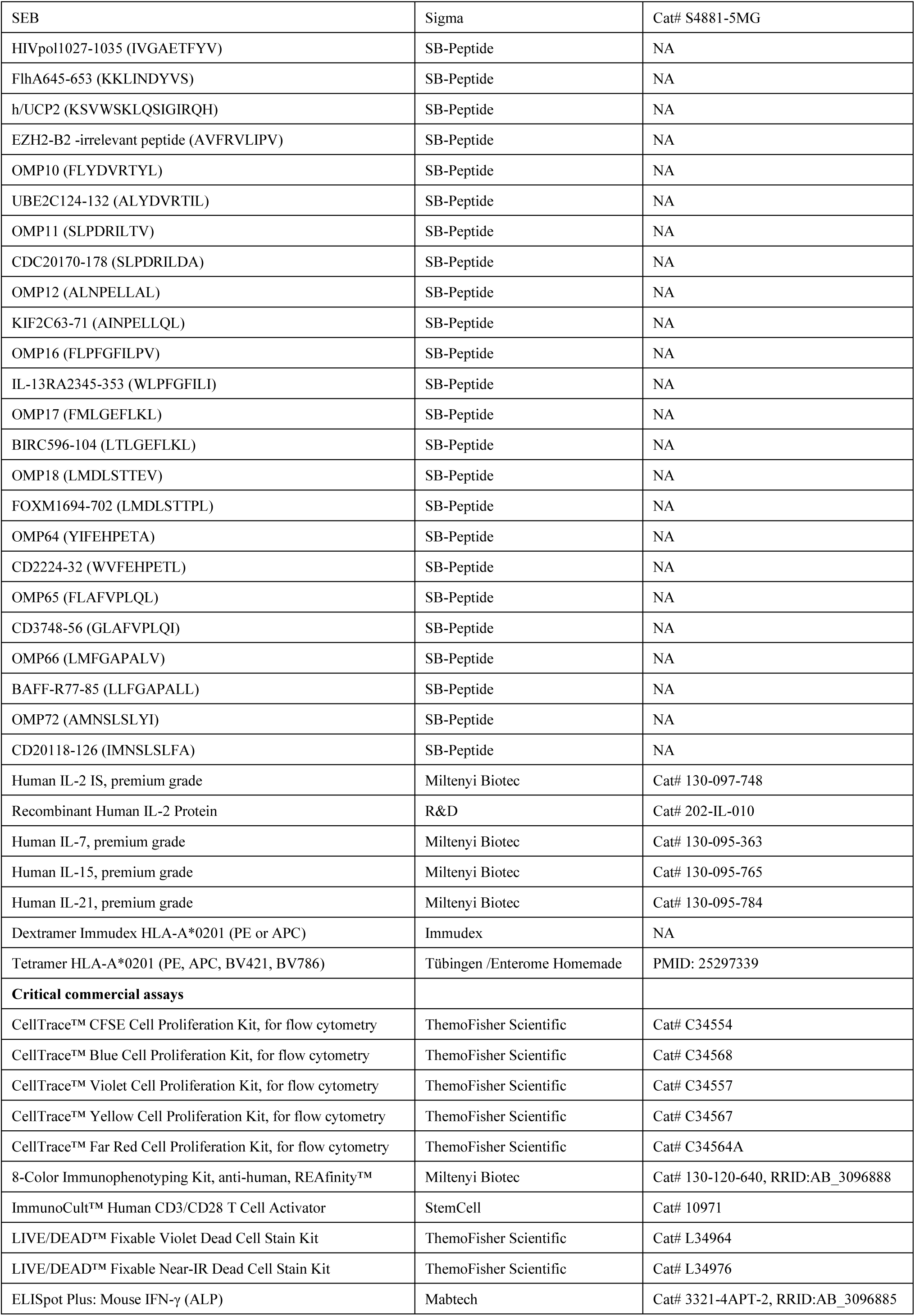
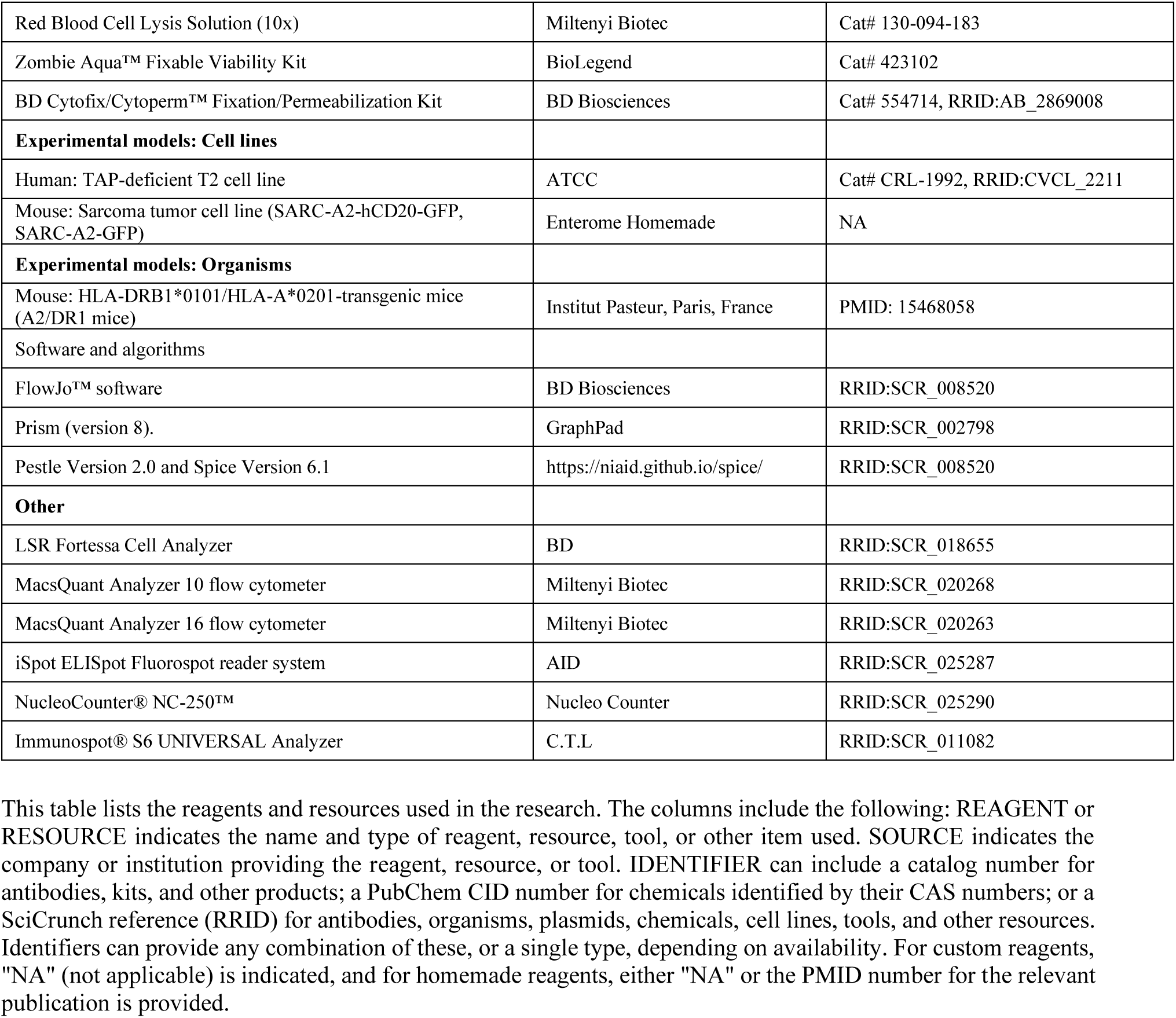

